# TrkB agonists prevent post-ischemic BDNF-TrkB mediated emergence of refractory neonatal seizures in CD-1 pups

**DOI:** 10.1101/2019.12.23.887190

**Authors:** P.A. Kipnis, B.J. Sullivan, B.M. Carter, S.D. Kadam

**Affiliations:** Neuroscience Laboratory, Hugo Moser Research Institute at Kennedy Krieger; Baltimore, MD 21205; Department of Neurology, Johns Hopkins University School of Medicine; Baltimore, MD 21205

## Abstract

Refractory neonatal seizures do not respond to first-line anti-seizure medications (ASMs) like phenobarbital (PB), a positive allosteric modulator for GABAA receptors, the most widely used ASM to treat neonatal seizures. GABAA receptor-mediated inhibition is dependent upon neuronal chloride regulation. The electroneutral cation-chloride transporter KCC2 mediates neuronal chloride extrusion; an age-dependent increase of KCC2 expression enables the shift of GABAergic signaling from depolarizing to hyperpolarizing. BDNF-TrkB activation following excitotoxic injury recruits downstream targets like PLCγ1, leading to KCC2 hypofunction. This study investigated the efficacy of partial and full TrkB agonists; LM22A-4 (LM), HIOC and Deoxygedunin (DG) respectively, on PB-refractory seizures, post-ischemic TrkB-pathway activation, and KCC2 membrane stability in a P7 CD-1 mouse model of refractory neonatal seizures. Anti-seizure efficacy was determined by quantifying seizure burdens with continuous video-EEG. LM rescued PB-refractory seizures in a sexually dimorphic manner. LM anti-seizure efficacy was associated with a significant reduction in the post-ischemic phosphorylation of TrkB at Y816, a site known to mediate post-ischemic KCC2 hypofunction via PLCγ1 activation. LM additionally rescued ischemia-induced pKCC2-S940 dephosphorylation preserving its membrane stability. HIOC and DG, two novel full TrkB agonists, also rescued PB-refractoriness and post-ischemic TrkB-PLCγ1 pathway activation. Additionally, chemogenetic inactivation of TrkB significantly reduced post-ischemic neonatal seizure burdens at P7. Developmental expression profiles of TrkB and KCC2 in naïve pups identified developmental differences that may underlie the sex-dependent variance in anti-seizure efficacy. These results support a novel role for the TrkB receptor in the emergence of age-dependent refractory neonatal seizures.

## Introduction

Excitotoxic injury has been shown to phosphorylate tyrosine receptor kinase B (TrkB) pathway signaling (*1–3*). TrkB is activated by its endogenous ligand, the neurotrophin brain-derived neurotropic factor (BDNF), and leads to the activation of multiple intracellular signaling cascades, including three major downstream signaling cascades: phospholipase Cγ1 (PLCγ1)/protein kinase C (PKC), mitogen-activated protein kinase (MAP/ERK kinase) and the phosphatidylinositol 3-kinase/Akt (*4*). We have previously demonstrated the post-ischemic activation of the TrkB-PLCγ1 pathway results in the hypofunction of the K-Cl co-transporter (KCC2) in a mouse model of acute neonatal ischemia associated with phenobarbital (PB)-refractory seizures (*5*).

The electroneutral cation-chloride transporter KCC2 is the primary neuronal Cl^-^ extruder and enables hyperpolarizing GABAergic inhibition in the brain. The residue Ser940 (S940) on the C-terminus of KCC2 is associated with its membrane stability and chloride extrusion capacity (*6, 7*). The developmental switch in GABAergic signaling from depolarizing to hyperpolarizing (*8*) is enabled by an age-dependent increase of KCC2 expression (*9*). In the neonatal period, KCC2 expression is low and GABA is depolarizing (*8–10*). In addition, KCC2 is susceptible to degradation following excitotoxic injury (*2, 11, 12*). In our characterized CD-1 mouse model of ischemic neonatal seizures, KCC2 underwent degradation and dephosphorylation of residue S940 (*5*). This rendered the ASM PB inefficacious (*5*), as PB is a positive allosteric modulator of GABAA receptors (*13*). Prevention of BDNF-TrkB mediated KCC2 hypofunction rescued PB-refractoriness in CD-1 pups (*5*). We hypothesized that the BDNF mimetic LM22A-4 (LM) (*14*) would interfere with the post-ischemic BDNF-TrkB signaling underlying the emergence of refractoriness.

This study utilized a characterized model of PB-refractory neonatal ischemic seizures at P7 and PB-responsive ischemic seizures at P10 (*2, 5, 15*). The efficacy of two graded doses (0.25mg/kg [LM] and 2.5mg/kg [LM2.5]) of the BDNF loop II mimetic, LM, was compared to the novel full TrkB agonists HIOC (*16*) and DG (*17*). HIOC is an *N*-acetylserotonin (NAS) derivative that exhibits more robust neurotrophic effects than NAS in a TrkB-dependent manner (*16, 18*). DG is a naturally occurring compound in the gedunin family that has shown robust neuroprotective properties in rats in a TrkB-dependent manner (*17*). The developmental profile of TrkB in neonatal brains is shown to decrease with age (*19–22*) and was also investigated here in the CD-1 strain. The role of TrkB receptor in neonatal seizure susceptibility was investigated using a chemogenetic mouse model.

## Materials and Methods

### Experimental Design

All experiments were done in compliance with the Johns Hopkins University Committee on the Ethics of Animal Experiments (Permit A3272-01) and approved by the Animal Care and Use Committee of Johns Hopkins. Litters of CD-1 pups were purchased from Charles River Laboratories, Inc. (Wilmington, MA). Pups with dams (litter size=10) were delivered on postnatal day 3 (P3) and were allowed to acclimate. Food and water were provided *ad libitum*. Doses for LM used in vivo and in vitro were determined from previously described *in vitro* pharmacokinetics (*14*). Pups of both sexes at P7 and P10 (see Table S1 for sample sizes) were intraperitoneally (IP; Fig. 1A) administered 0.25mg/kg LM dissolved in isotonic phosphate-buffered saline (PBS) as two treatment groups: 1. one dose 2h before ligation (Pre LM), 2. immediately following ligation (Post LM). After 1h of baseline EEG recording, pups were given a loading dose of PB (25mg/kg, IP). Doses for HIOC used in vivo and in vitro were determined from previously described in vitro pharmacokinetics (*16*). Pups of both sexes at P7 were IP administered the full agonist selective for TrkB 5mg/kg N-[2-(5-hydroxy-1H-indol-3-yl) ethyl]-2-oxopiperidine-3-carboxamide (HIOC) dissolved in 95/5% isotonic PBS/DMSO as two treatment groups: 1. one dose 2h before ligation (Pre HIOC), 2. immediately following ligation (Post HIOC). After 1h of baseline EEG recording, pups were given a loading dose of PB (25mg/kg, IP). Pups of both sexes at P7 were IP administered 5mg/kg the full agonist selective for TrkB, deoxygedunin (DG), dissolved in 95/5% isotonic PBS/DMSO as two treatment groups: 1. one dose 2h before ligation (Pre DG), 2. immediately following ligation (Post DG). After 1h of baseline EEG recording, pups were given a loading dose of PB (25mg/kg, IP). During the course of the experiments, DG became commercially unavailable, which is reflected for the smaller sample size for the DG treated groups. Our group has previously published results showing that the 5% DMSO used as a vehicle for drug administration did not have any anti-seizure effect and did not alter baseline seizure burdens (*2*).

**Fig. 1.**
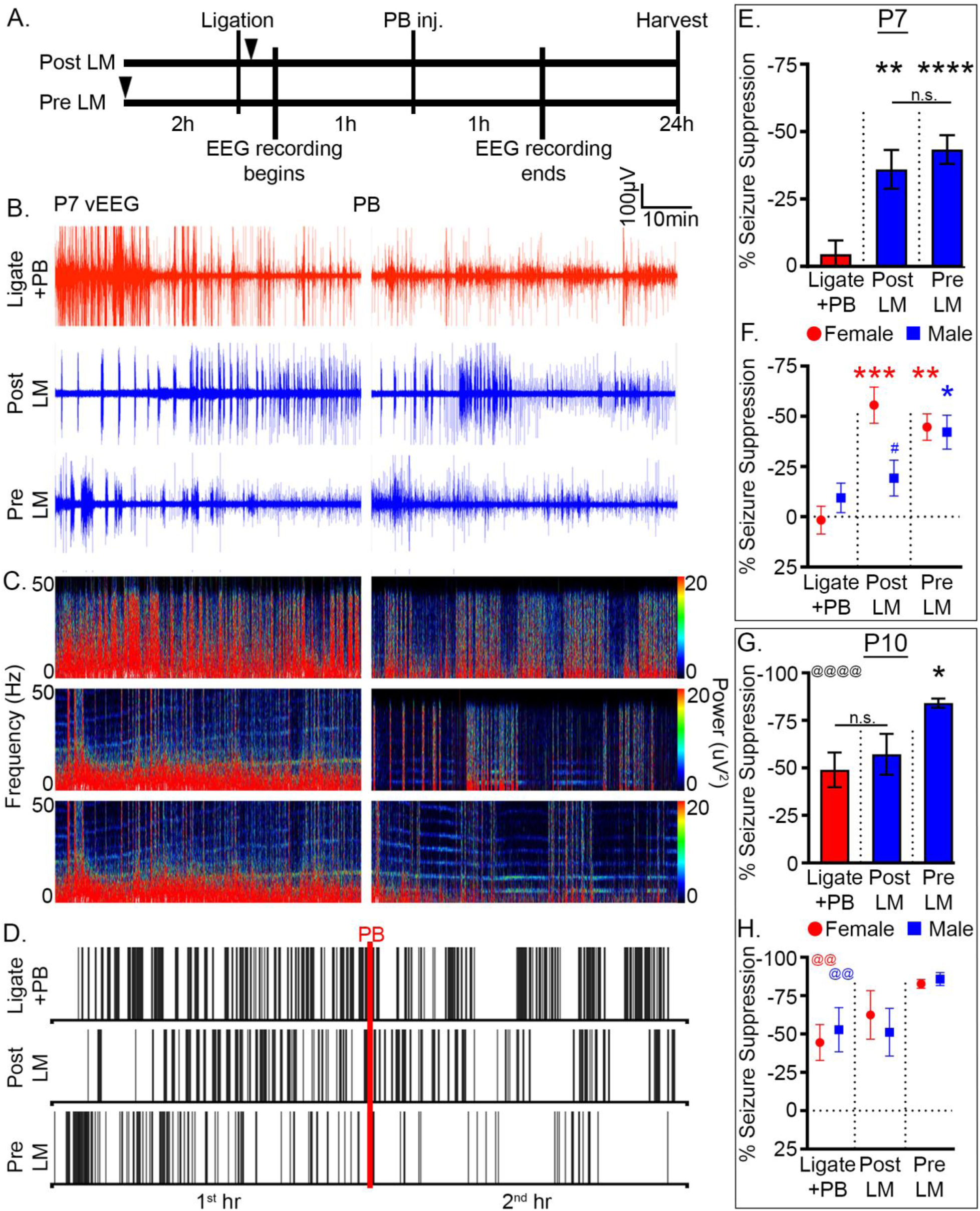
LM significantly rescued PB-refractoriness. (A) Experimental paradigm to evaluate LM efficacy in a mouse model of ischemic neonatal seizures. Pups were randomly assigned to treatment groups (see Table S1 for sample sizes). Black arrowheads indicate time point of LM intervention. **(B)** Representative EEG traces, **(C)** power spectrograms (0-50Hz), and **(D)** raster plots showing significantly rescued PB-refractoriness in Post and Pre LM pups. Red line indicates time point of PB administration. **(E)** EEG Percent seizure suppression for Ligate+PB, Post LM, and Pre LM treated P7 pups. ** p<0.01 (Post LM vs. Ligate+PB), **** p<0.0001 (Pre LM vs. Ligate+PB) by one-way ANOVA. **(F)** EEG Percent seizure suppression by sex for Ligate+PB, Post LM, and Pre LM treated P7 pups. * p<0.05 (Male Pre LM vs. Male Ligate+PB), ** p<0.01 (Female Pre LM vs. Female Ligate+PB), *** p<0.001 (Female Post LM vs. Female Ligate+PB) by two-way ANOVA. # indicated within-group sex differences; # p<0.05 by two-tailed *t* test. **(G)** EEG Percent seizure suppression for Ligate+PB, Post LM, and Pre LM treated P10 pups. * p<0.01 (Pre LM vs. Ligate+PB) by one-way ANOVA. @ indicates difference between P7 and P10; @@@@ p<0.0001 by two-tailed *t* test. **(H)** EEG Percent seizure suppression by sex for Ligate+PB, Post LM, and Pre LM treated P10 pups. @ indicates sex differences between P7 and P10; @@ p<0.01 by two-tailed *t* test.

To run analysis for drug efficacies, data for naïve male and female pups were pooled across all treatment groups for P7. The Ligate+PB group was pooled from the LM, LM 2.5mg/kg, and HIOC treatment litter mates. Experiments for DG were performed as an additional positive control and the Ligate+PB group was pooled for LM, LM 2.5mg/kg, and DG treatment litter mates (see Table S1 for sample size details).

### Surgery Protocol for Carotid Ligation and EEG Electrode Implantation

At P7 or P10, pups underwent permanent unilateral ligation of the right common carotid artery without transection using 6-0 surgisilk (Fine Science Tools, USA) under isofluorane anesthesia (Henry Schein, USA). The skin was closed with 6-0 monofilament nylon (Covidien) and lidocaine was applied as a local anesthetic. Pups were then implanted with three subdermal EEG electrodes (SWE-L25, Ives EEG Solutions, USA): 1 recording and 1 reference overlying the left/right parietal cortex, and 1 ground overlying the rostrum (*15, 23*). The electrodes were fixed in place using cyanoacrylate adhesive (KrazyGlue). Pups were tethered to a preamplifier inside the recording chamber and allowed to recover from anesthesia. vEEG was recorded continuously for 2h in a chamber maintained at 37°C with isothermal pads. At the end of recording, the electrodes were removed and pups were returned to the dam.

### EEG Recordings and Analyses

EEG recordings were acquired using Sirenia Acquisition (v1.6.4, Pinnacle Technology, Inc., USA) with synchronized video recording. Data were recorded with a 400Hz sampling rate that had a preamplifier gain of 100, and 0.5-50Hz low-pass filter to remove ambient noise. The EEG data were then binned into 10s epochs for manual scoring. Seizures were defined as electrographic events consisting of rhythmic spikes of high amplitude, diffuse peak frequency of ≥7-8Hz lasting ≥6 seconds, similar to previous studies (*5, 23*). Similarly to previously published protocols in this model, short-duration burst activity lasting <6s was not included in seizure burden calculations. Seizure suppression was calculated as:

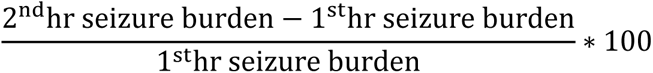

### EEG Power Analysis

Four P7 pups of each sex in each treatment group were randomly chosen for EEG power analysis. EEG power was obtained with Sirenia Sleep (v1.7.10, Pinnacle Technology Inc., USA). EEG spectral power from 0.5-50Hz was acquired for each 10s epoch of recording after automated fast Fourier transformation. Data from EEG artifacts was excluded from these analyses. Total EEG power was calculated as follows:

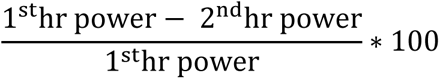

### P7 ligation in C57BL/6 Knockin TrkB^F616A^ Pups

*TrkB^F616A^* breeding pairs (JAX stock #022363, developed by the Dr. David Ginty lab, (*24*)) were obtained courtesy of Dr. Richard Huganir (Johns Hopkins University). To investigate the effect of chemogenetically induced deficits in neonatal TrkB signaling in vivo, P7 pups with *TrkB^F616A^* knockin alleles (F616A^+/+^) (*24*) received the kinase inhibitor 1NMPP1 from P0 to P7 via transmammary route with the dam receiving the chemogenetic agent in her drinking water (10% TWEEN-20 and 80uM 1NMPP1 in drinking water). P7 WT^-/-^ and F616A^+/+^ littermates with 1NMPP1 treatment underwent unilateral carotid ligation and subsequent qEEG at P7 as previously described.

### Western Blotting

24h after ligation all animals were anesthetized with 0.1mL of 90mg/mL chloral hydrate (IP). Pups were then transcardially perfused with ice cold PBS followed by 1 mL of 1x HALT protease inhibitor cocktail in PBS (ThermoFisher #78430, USA). The whole brains were removed, the cerebellum was discarded, and the left and right hemispheres were separated. For the developmental series, brains were further micro-dissected into cortex, hippocampus, and deep gray matter and stored at -80°C until further processing. Homogenized brain tissue was suspended in T-PER cell lysis buffer containing 1x HALT protease inhibitor cocktail. Protein concentration was measured using the Bradford protein assay at 570nm. 25µg of total protein (60µg for PLCγ1) in 20µL was run on 4-20% tris-glycine gels (ThermoFisher #XP04205BOX, USA) for 120 minutes at 130V. Gels were transferred overnight at 20V onto nitrocellulose membranes. Membranes were blocked in Rockland Blocking Buffer for 1h (Rockland #MB-070, USA). Membranes underwent 6h incubation with primary antibodies: mouse α KCC2 (1:1000, Millipore; #07-432), rabbit α phospho-KCC2(S940) (1:1000 Aviva Systems Biology; #OAPC00188), mouse α TrkB (1:1000, BD Biosciences; #610102), rabbit α phospho-TrkB(T816) (1:500, Millipore; #ABN1381), mouse α PLCγ1 (1:1000, Thermo Scientific; #LF-MA0050), rabbit α phospho-PLCγ1(T783) (1:1000, Cell Signaling Technology; #2821S), rabbit α Erk1/2 (1:1000, Cell Signaling Technology; #4695), rabbit α phospho-Erk1/2 (1:1000, Cell Signaling Technology; #4377), and mouse α β-actin (1:10,000, Li-Cor; 926-42212). Membranes were then incubated with fluorescent secondary antibodies (1:5000, Li-Cor 926-68020 and 925-32211, USA; for antibody RRIDs, see Table S2). Blots were visualized on the Odyssey infrared imagining system 2.1 (Li-Cor Biosciences, USA). Optical density for each protein band was normalized to β-actin in the same lane.

### Surface Protein Separation by Ultracentrifugation

1mm coronal slices were obtained from P7 mouse brains and were allowed to recover for 45min at 34°C with oxygenation 95/5% O2/CO2. After recovery, slices were incubated with TrkB agonists LM22A-4 and HIOC at 34°C with oxygenation. Slices were placed in cell lysis buffer TPER with HALT protease and phosphatase inhibitors and homogenized with sonicator. After 30min incubation on ice, protein lysates were ultracentrifuged at 70K RPM and supernatant was collected as cytosolic sample. Pellets were resuspended and ultracentrifuged, with supernatant discarded as wash component. Pellets were resuspended and collected as membrane component. Membrane and cytosolic components underwent Bradford analysis and Western blotting for protein quantification.

### Statistical Analyses

All statistical analyses were performed in Prism 7 (Graphpad, USA). Percent seizure suppression by sex, seizure burdens, ictal events, ictal durations, and protein expression levels quantified from P5 to P21 were analyzed using two-way ANOVA and post-hoc corrections were made using Tukey’s test. Percent seizure suppression and all western blot data at P7 and P10 for drug efficacies were analyzed using one-way ANOVA and post-hoc corrections were made using Sidak’s test. For comparisons between ipsi- and contralateral hemispheres within groups at P7 and P10, as well as comparisons of baseline seizure burdens between developmental ages P7 and P10, two-tailed t-tests were performed. Significance of correlations between percent EEG spectral power suppression and percent seizure burden suppression across treatment groups were performed using Spearman’s two-tailed nonparametric test. An alpha of p<0.05 was considered significant. All data represent the mean ± 1 standard error of the mean (SEM).

## Results

### LM rescued neonatal PB-refractory seizures in a sex-dependent manner

Ischemia-induced seizures in P7 CD-1 pups are PB-refractory (*15*). Previously it has been shown that ANA12, a novel small-molecule TrkB antagonist, significantly rescued PB-refractoriness by blocking BDNF-TrkB pathway activation in a dose dependent manner (*2, 5*). To evaluate the efficacy of LM22A-4, a small-molecule TrkB partial agonist on rescuing PB-refractoriness, P7 pups were either given LM immediately after ligation (Post LM) or 2h before ligation (Pre LM) per the experimental paradigm (Fig. 1A). Continuous 2h vEEG/EMG recordings were used to identify and quantify post-ischemic electrographic seizure burdens (Fig. 1B-D). The seizure burden in the Ligate+PB group remained unchanged following PB administration, indicating PB-refractoriness. The Post and Pre LM groups both showed significant rescue of PB-refractoriness (Fig. 1B-E). Clustering of ictal events was noted in the Ligate+PB raster plot following PB injection (i.e. Ligate+PB 2^nd^ h vs. 1^st^ h) without overall reduction in seizure burdens (Fig. 1D). Both the Post and Pre LM groups also showed similar clustering of ictal events, although this was associated with a concomitant significant reduction in overall seizure burdens following PB administration (Fig. 1D and E). When quantified as percent seizure suppression over the 1^st^ hour baseline, intervention with LM+PB significantly increased seizure suppression in P7 seizing pups (Fig 1E), in contrast to intervention with only PB which failed to show seizure suppression (-4.46±5.34%).

Neonatal seizure susceptibility and ASM efficacy has been shown to be sexually dimorphic (*5, 15, 23, 25, 26*), therefore sex as a biological variable was investigated. At P7, both sexes were PB-refractory. Female pups were significantly responsive to LM intervention in both the Post and Pre LM groups, whereas male pups only responded in the Pre LM group (Fig. 1F). Furthermore, percent seizure suppression in female pups was not significantly different between the Post and Pre LM groups, demonstrating that pre-ischemic LM intervention did not provide females any additional benefit in the rescue of PB-refractoriness. However, female pups in the Post LM group showed significantly greater percent seizure suppression than male pups in the Post LM group, highlighting important sex differences in the efficacy of LM at P7.

As previously characterized for the model, neonatal ischemic seizures at P10 were PB-responsive in both sexes (*15*) (Fig. 1G, H), supporting a developmental influence on PB-efficacy. At P10, only the Pre LM group showed significant improvement in PB-efficacy (Fig. 1G). In contrast to P7, no sexual dimorphism was noted for either the Post or Pre LM treatment groups at P10 (Fig. 1H), highlighting the role of developmental age in sexual dimorphism. In summary, LM intervention significantly rescued PB-refractoriness at P7 and improved PB-efficacy at P10.

### No dose-dependent effect for a graded dose of LM

To evaluate the dose-dependent efficacy of LM, a ten-fold higher dose (0.25 vs. 2.5mg/kg) was evaluated for PB-refractory seizures at P7. The Post LM2.5 group significantly suppressed PB-refractory seizures by 42.77±9.98%, similar to but not significantly better than the Post LM 0.25mg/kg dose (two-tailed *t* test, p=0.6375). In contrast, the Pre LM2.5 group failed to significantly rescue PB-refractory seizures (18.15±16% seizure suppression, Fig. S1A). In contrast to the 0.25mg/kg dose, there were no significant differences in seizure suppression between sexes in the Post and Pre LM2.5 groups (Fig. S1B). The significant seizure suppression in both the Post LM and Post LM 2.5 groups indicated a nonlinear dose-response curve (*27*).

To evaluate the effect of TrkB agonists LM and HIOC on plasma membrane expression of TrkB and KCC2, naïve P7 mouse pup brain slices were incubated with either 0.75mM or 7.5mM graded doses of LM, or a 1.7mM dose of HIOC. These in vitro doses mimicked the in vivo doses used to study anti-seizure efficacies (.25mg/kg, 2.5mg/kg, and 5mg/kg respectively). Incubation of naïve P7 brain slices with the TrkB agonists showed no significant increase in pTrkB-Y816 and KCC2 expression at the plasma membrane (Fig S1C, D). Similarly the ratio of pTrkB-Y816 to total TrkB at the membrane failed to show a significant dose-dependent increase with either LM or HIOC (Fig. S1E). The S940/KCC2 ratios at the membrane were also not significantly modulated by LM or HIOC (Fig. S1F). These in vitro findings indicate that the rescue of KCC2 expression by LM and HIOC is specific to the post-ischemic activation of the BDNF-TrkB pathway in seizing pups. In the absence of the ischemic injury, the TrkB agonists did not significantly modulate either TrkB or KCC2 insertion at the membrane.

### Effect on frequency and duration of neonatal ischemic seizures

To investigate the seizure semiology, seizure burden was evaluated as the total amount of time spent seizing on EEG. Baseline seizure burdens (i.e.; represented by the seizures during the 1^st^ hour of EEG recording) for each treatment group in this study were not significantly different from each other at P7 (Fig. 2A) and P10 (Fig. 2B). At P7, the Post and Pre LM groups demonstrated significant reductions in seizure burdens during the 2^nd^ hour following PB administration (Fig. 2A). In contrast, the Ligate+PB group failed to reduce seizure burdens in the 2^nd^ hour, demonstrating PB-refractoriness. At P10 for the same ischemic insult, baseline seizure burdens were significantly lower than at P7 (Fig. 2B vs. 2A), as previously characterized for the model (*2, 5, 15*). Both the Post and Pre LM groups showed significant reduction in seizure burdens following PB administration, similar to the Ligate+PB group.

**Fig. 2.**
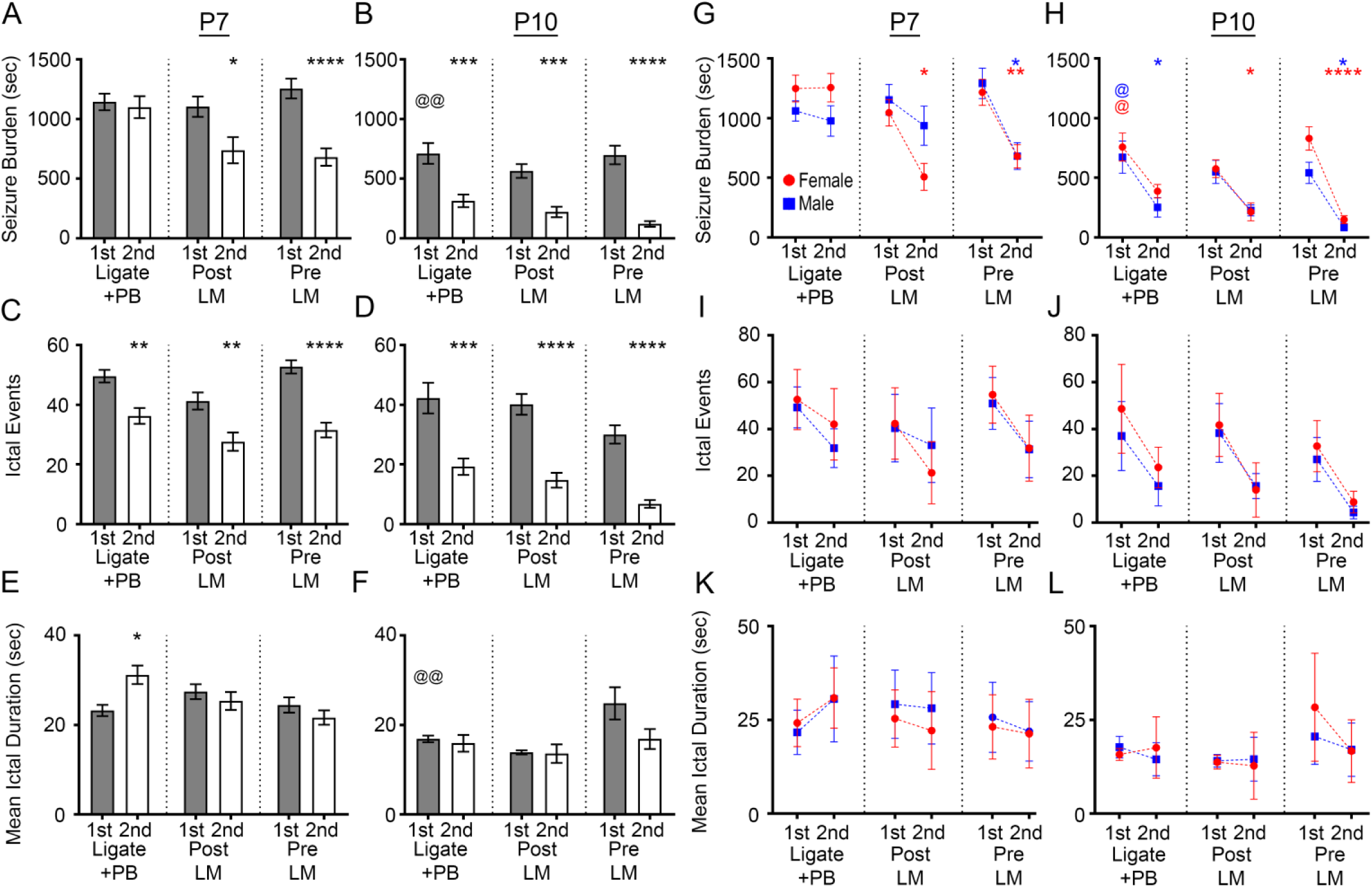
LM efficacy was sexually dimorphic and driven by significant reduction of ictal events, but not ictal durations at both P7 and P10. (A) EEG seizure burdens for Ligate+PB, Post LM, and Pre LM treated P7 pups. * p<0.05 (2^nd^ Post LM vs. 1^st^ Post LM), **** p<0.0001 (2^nd^ Pre LM vs. 1^st^ Pre LM) by two-way ANOVA. **(B)** EEG seizure burdens for Ligate+PB, Post LM, and Pre LM treated P10 pups. *** p<0.001 (2^nd^ Ligate+PB vs. 1^st^ Ligate+PB, and 2^nd^ Post LM vs. 1^st^ Post LM), **** p<0.0001 (2^nd^ Pre LM vs. 1^st^ Pre LM) by two-way ANOVA. @ indicates differences between P7 and P10; @@ p<0.01 by two-tailed *t* test. **(C)** EEG ictal events for Ligate+PB, Post LM, and Pre LM treated P7 pups. ** p<0.001 (2^nd^ Ligate+PB vs. 1^st^ Ligate+PB, and 2^nd^ Post LM vs. 1^st^ Post LM), **** p<0.0001 (2^nd^ Pre LM vs. 1^st^ Pre LM) by two-way ANOVA. **(D)** EEG ictal events for Ligate+PB, Post LM, and Pre LM treated P10 pups. *** p<0.001 (2^nd^ Ligate+PB vs. 1^st^ Ligate+PB), **** p<0.0001 (2^nd^ Post LM vs. 1^st^ Post LM, and 2^nd^ Pre LM vs. 1^st^ Pre LM) by two-way ANOVA. **(E)** EEG mean ictal durations for Ligate+PB, Post LM, and Pre LM treated P7 pups. * p<0.05 (2^nd^ Ligate+PB vs. 1^st^ Ligate+PB) by two-way ANOVA. **(F)** EEG mean ictal durations for Ligate+PB, Post LM, and Pre LM treated P10 pups. **(G)** EEG seizure burdens by sex for Ligate+PB, Post LM, and Pre LM treated P7 pups. * p<0.05 (2^nd^ Female Post LM vs. 1^st^ Female Post LM, and 2^nd^ Male Pre LM vs. 1^st^ Male Pre LM), ** p<0.01 (2^nd^ Female Pre LM vs. 1^st^ Female Pre LM) by two-way ANOVA. **(H)** EEG seizure burdens by sex for Ligate+PB, Post LM, and Pre LM treated P10 pups. * p<0.05 (2^nd^ Male Ligate+PB vs. 1^st^ Male Ligate+PB, 2^nd^ Female Post LM vs. 1^st^ Female Post LM, and 2^nd^ Male Pre LM vs. 1^st^ Male Pre LM), **** p<0.0001 (2^nd^ Female Pre LM vs. 1^st^ Female Pre LM) by two-way ANOVA. @ p<0.05 by two-tailed *t* test. **(I)** EEG ictal events by sex for Ligate+PB, Post LM, and Pre LM treated P7 and **(J)** P10 pups. **(K)** EEG mean ictal durations by sex for Ligate+PB, Post LM, and Pre LM treated P7 and **(L)** P10 pups.

Analysis of the number of ictal events (Fig. 2C and D) in all treatment groups revealed that the significant seizure suppression (Fig. 1) with LM intervention was driven by significant reductions in the number of ictal events at both P7 and P10 (Fig. 2C and D). PB-refractoriness in the Ligate+PB group at P7 was driven by a significant increase in ictal durations (Fig. 2E). In contrast, ictal durations during the 2^nd^ hour in the Post and Pre LM groups were not significantly different at both P7 and P10 (Fig. 2E and F). P7 female pups in both the Post and Pre LM groups had significantly lower 2^nd^ hour seizure burdens than their respective 1^st^ hour (Fig. 2G). In contrast, males only in the Pre LM group had significantly lower 2^nd^ hour seizure burdens when compared to their respective 1^st^ hour at P7.

At P10, the 1^st^ hour seizure burdens for both male and female pups in the Ligate+PB group had significantly lower 1^st^ hour seizure burdens than their respective P7 counterparts (Fig. 2H). In contrast to P7, the P10 Ligate+PB group demonstrated differences between sexes as only males had significantly lower seizure burdens in the 2^nd^ hour than their respective 1^st^ hour. The females in the Post LM group at P10 had significantly lower seizure burdens in the 2^nd^ hour than their respective 1^st^ hour, similar to P7. In contrast, both P10 males and females in the Pre LM group had significantly lower 2^nd^ hour seizures burdens than their respective 1^st^ hour, similar to Pre LM at P7. The sex-dependent differences in ictal events and durations at both ages were not significant (Fig. 2I-L).

### EEG power was not predictive of acute ASM efficacy

EEG power has been used as a proxy to determine seizure burden on acute induced seizures (*28, 29*). EEG power suppression was examined to evaluate efficacy of LM intervention (for example 10min EEG seizure trace see S2A-D). The Ligate+PB group showed similar percent EEG power suppression to the LM-treated groups (Fig. S2E), which was driven by reductions in the 2^nd^ hour EEG powers in all treatment groups. Therefore, EEG spectral power evaluated in a subset of LM-treated P7 pups failed to estimate accurate seizure burdens both in the Ligate+PB and LM-treated groups (Fig. S2F), indicating the unreliability of EEG power to detect accurate seizure burdens. Overall EEG power diminishes with the occurrence of repeated ischemic seizures (*30*). This phenomenon has also been reported for clinical EEGs (*31*). The correlation between percent EEG power suppression and percent seizure suppression showed that quantification of EEG power alone could not accurately measure seizure burdens (Fig. S2G), similar to previous reports (*5, 23*).

### TrkB inactivation facilitates post-ischemic amelioration of seizure susceptibility at P7

To investigate the role of post-ischemic TrkB-pathway activation at P7 in vivo, TrkB activation was chemogenetically inhibited by 1NMPP1 from P0-P7 in *C57BL/6 Knockin TrkB^F616A^* (F616A^+/+^) pups. Following unilateral carotid ligation, WT^-/-^ pups administered 1NMPP1 showed significantly higher 1^st^ h seizure burdens than F616A^+/+^ pups administered 1NMPP1 (Fig. 3A-D) demonstrating that the chemogenetic inactivation of TrkB significantly reduced the post-ischemic seizure susceptibility in P7 neonatal pups. EEG seizure burdens were significantly lower in the F616A^+/+^ pups in the 1^st^ h following ischemia. EEG seizure burdens evaluated by sex, found no sex dependent differences. PB-administration at the end if 1h in WT^-/-^ *C57BL/6* pups (Fig 3D) was efficacious. These results support the established importance of genetic background on ischemia and seizures as phenotypic severity is strain dependent (*32–34*). Specifically, the CD-1 strain shows phenotype severity and emergence of refractory neonatal seizures with added translational value in comparison to C57BL/6 strain. Overall, these results demonstrate that post-ischemic activation of the BDNF-TrkB signaling cascade plays a crucial role in neonatal seizure susceptibility.

**Fig. 3.**
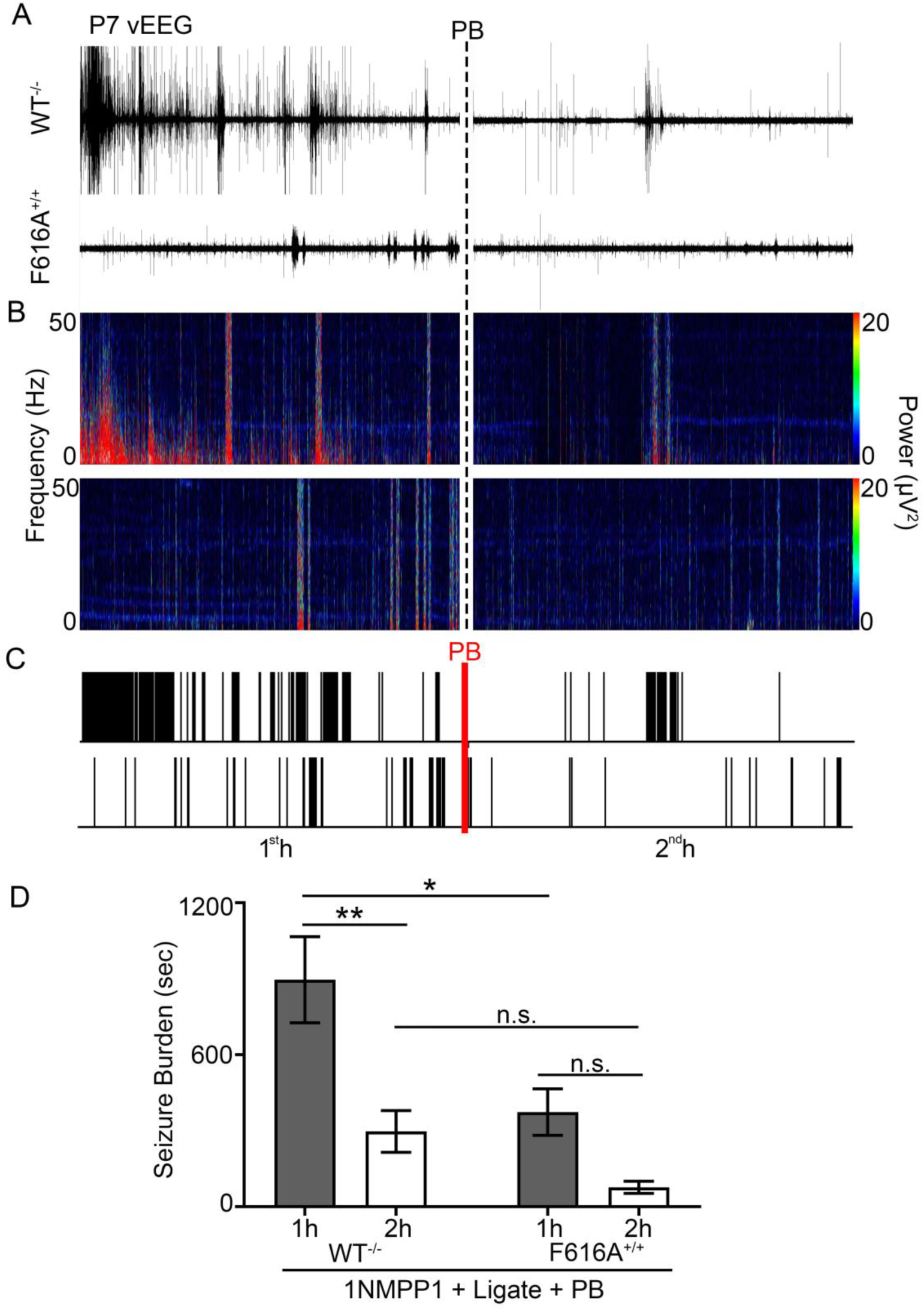
Post-ischemic TrkB activation underlies neonatal seizure susceptibility. (A) Representative EEG traces in WT^-/-^ and mutant F616A^+/+^ pups, **(B)** power spectrograms (0-50Hz), and **(C)** raster plots showed significantly lower 1^st^ h post-ischemic EEG seizure burdens in F616A^+/+^ pups. Dotted black and red lines indicate time point of PB administration. **(D)** EEG seizure burdens during 1^st^ and 2^nd^ h post-ligation in WT^-/-^ and F616A^+/+^ pups. * p<0.05, ** p<0.01 by two-way ANOVA.

### Post-ischemic TrkB-PLCγ1 pathway activation at P7 was rescued by LM intervention

Ischemic insults are known to induce BDNF-TrkB pathway activation (*5, 35*), and phosphorylation of Y816 on TrkB activates the PLCγ1 pathway, which has been implicated both acutely (*5*) and chronically in epileptogenesis (*36*). 24h post-ischemia, pups in the Ligate+PB group showed significant increase of TrkB and pTrkB-Y816 expression ipsi- and contralateral to ischemic insult (Fig. 4A-C), indicating global TrkB activation in the unilateral model. With the unilateral ischemia model used in this study, ispi- vs. contralateral-hemispheric differences in protein expression were also analyzed. The Post LM group showed attenuated TrkB expression ipsilateral to ischemic insult (Fig. 4A). The pTrkB-Y816 / TrkB ratio of the Ligate+PB group was significantly lower ipsilateral to insult (Fig. S3A). The Post and Pre LM groups both significantly rescued post-ischemic TrkB-pathway activation. The ratios of pTrkB-Y816 to total TrkB showed no significant differences between treatment groups (Fig. S3A). Post-ischemic TrkB activation was analyzed by sex, and no significant sex-dependent differences were found.

**Fig. 4.**
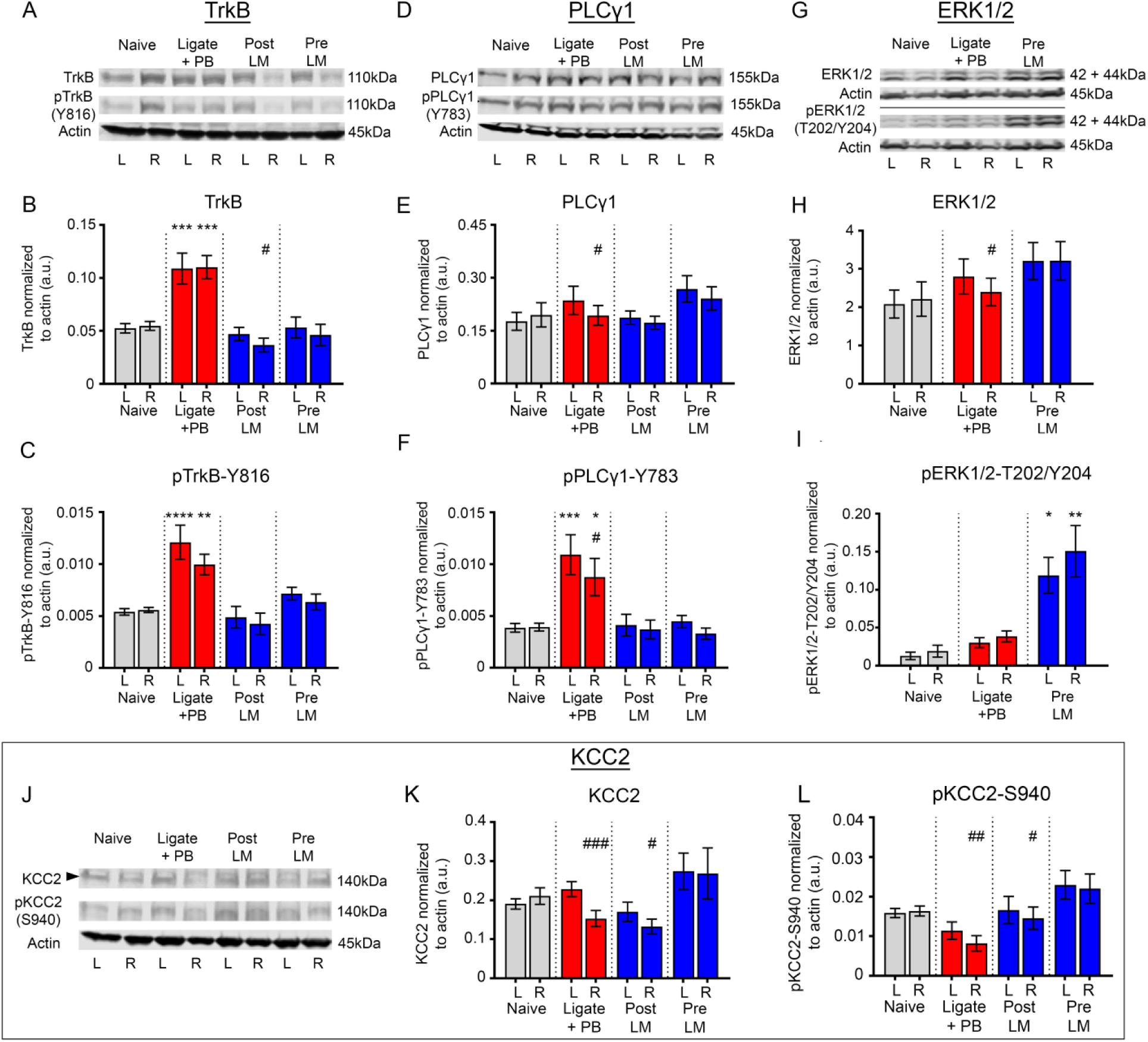
LM rescued post-ischemic TrkB-PLCγ1 pathway activation, activated the TrkB-ERK1/2 pathway, and rescued ipsilateral KCC2 degradation. All proteins of interest were normalized to housekeeping protein β-actin. Phospho-proteins were also normalized to their respective total protein (see Figure S3). * indicate differences between treatment group and naïve controls; # indicated differences between contralateral and ipsilateral hemispheres within groups. **(A)** Representative Western blots showing TrkB and pTrkB-Y816 expression for all treatment groups. **(B)** Contralateral (L) and ipsilateral (R) TrkB expression 24h after ischemic insult for all treatment groups. *** p<0.001 by one-way ANOVA. # p<0.05 by two-tailed *t* test. **(C)** Contralateral (L) and ipsilateral (R) pTrkB-Y816 expression 24h after ischemic insult for all treatment groups. ** p<0.01, **** p<0.0001 by one-way ANOVA. **(D)** Representative Western blots showing PLCγ1 and pPLCγ1-Y783 expression for all treatment groups. **(E)** Contralateral (L) and ipsilateral (R) PLCγ1 expression 24h after ischemic insult for all treatment groups. # p<0.05 by two-tailed *t* test. **(F)** Contralateral (L) and ipsilateral (R) pPLCγ1-Y783 expression 24h after ischemic insult for all treatment groups. * p<0.05, *** p<0.001 by one-way ANOVA. # p<0.05 by two-tailed *t* test. **(G)** Representative Western blots showing ERK1/2 and pERK1/2- T202/Y204 expression for all treatment groups. **(H)** Contralateral (L) and ipsilateral (R) ERK1/2 expression 24h after ischemic insult for all treatment groups. # p<0.05 by two-tailed *t* test. **(I)** Contralateral (L) and ipsilateral (R) pERK1/2-T202/Y204 expression 24h after ischemic insult for all treatment groups. * p<0.05, ** p<0.01 by one-way ANOVA. **(J)** Representative Western blots showing KCC2 and pKCC2-S940 expression for all treatment groups. **(K)** Contralateral (L) and ipsilateral (R) KCC2 expression 24h after ischemic insult for all treatment groups. # p<0.05, ### p<0.001 by two-tailed *t* test. **(L)** Contralateral (L) and ipsilateral (R) pKCC2-S940 expression 24h after ischemic insult for all treatment groups. # p<0.05, ## p<0.01 by two-tailed *t* test.

Total PLCγ1 expression was not significantly modulated by ischemic insult or Post and Pre LM intervention (Fig. 4D and E). In contrast, pPLCγ1-Y783 expression was significantly higher both ipsi- and contralateral to ischemic insult, which was rescued both in Post LM and Pre LM groups (Fig. 4D and F). PLCγ1 and pPLCγ1-Y783 expression levels were also significantly lower ipsilateral to insult (Fig. 4D-F). The ratio of pPLCγ1-Y783 to total PLCγ1 was not significantly different between treatment groups (Fig. S3B).

TrkB pathway activation is also known to activate downstream ERK1/2 signaling (*37*). At P7, Pre LM was the most efficacious treatment paradigm in reducing EEG seizure burdens when quantified as percent seizure suppression (Fig 1E) for both sexes. Therefore, ERK1/2 activation was investigated in the Pre LM group. The TrkB-ERK1/2 pathway was not significantly activated by ischemic insult and was not influenced by LM intervention. However, ERK1/2 expression was significantly lower ipsilateral to insult in the Ligate+PB group (Fig. 4G and H). Furthermore, the Pre LM group showed significant activation of pERK1/2-T202/Y204 (Fig 4I). These data indicate the BDNF loop II mimetic differentially activated the TrkB-ERK1/2 pathway while simultaneously blocking ischemia-induced activation of the TrkB-PLCγ1 pathway. The ratio of pERK1/2-T202/Y204 to total ERK1/2 was significantly higher in the Pre LM group both ipsi- and contralateral to insult (Fig. S3C).

### LM rescued post-ischemic KCC2 and pKCC2-S940 degradation at P7

Post-ischemic TrkB-PLCγ1 pathway activation and seizures lead to ipsilateral KCC2 degradation in this model of unilateral ischemic insult (*2, 5*). The effect of TrkB-PLCγ1 pathway activation on KCC2 expression was evaluated 24h post-ischemia in LM intervention groups. The Post and Pre LM treatment group significantly rescued the ipsilateral KCC2 degradation seen in the Ligate+PB group (Fig. 4J and K). pKCC2-S940 is associated with KCC2 stability on the plasma membrane and thus its functionality as a Cl^-^ extruder (*6*). Similar to KCC2, the ipsilateral pKCC2-S940 dephosphorylation in the Ligate+PB group was significantly rescued in the Post and Pre LM Groups (Fig. 4L). The ratio of pKCC2-S940 to total KCC2 was not significantly different between treatment groups (Fig. S3D). In summary, LM intervention rescued post-ischemic BDNF-TrkB-PLCγ1 pathway activation, thus preventing KCC2 endocytosis and subsequent hypofunction.

### Post-ischemic TrkB-PLCγ1 pathway activation was not evident in ischemic P10 pups

In contrast to P7 pups, no increase in pTrkB-Y816 expression was detected in any treatment group at P10 (Fig. S4A-C). The Pre LM group showed significantly higher ratios of pTrkB-Y816 to TrkB ipsi- and contralateral to insult (Fig. S4D). PLCγ1 expression was not different between treated and untreated pups, though Post and Pre LM pups showed ipsilateral downregulation of pPLCγ1-Y783 (Fig. S4E-G). Ratios of pPLCγ1-Y784 to total PLCγ1 were not affected by LM intervention (Fig. S4H). No significant KCC2 degradation was detected in the Ligate+PB group (Fig. S4I and J) following ischemia, though Post LM pups showed lower ipsilateral pKCC2-S940 expression (Fig. S4K, L). In summary, LM intervention showed mild activation of the TrkB pathway, but KCC2 levels were not significantly modulated at P10 when ischemic seizures were responsive to PB.

### TrkB agonists HIOC and DG rescued refractory ischemic seizures at P7 similar to LM

The efficacy of two full TrkB agonists HIOC (*16*) and DG (*17*) was investigated to determine if the anti-seizure efficacy of LM was TrkB site-specific. At P7, HIOC demonstrated significant seizure suppression in both the Post and Pre HIOC groups (Fig. 5A) indicating that LM anti-seizure efficacy was not loop II site-specific. Male pups in the Pre HIOC group demonstrated significant seizure suppression compared to male Ligate+PB pups, in contrast to female pups (Fig. 5B). No significant sex differences were noted in the Post HIOC group. The 2^nd^ hour seizure burden in the Pre HIOC group was significantly lower than its 1^st^ hour baseline (Fig. 5C), in contrast to the Post HIOC and Ligate+PB groups.

**Fig. 5.**
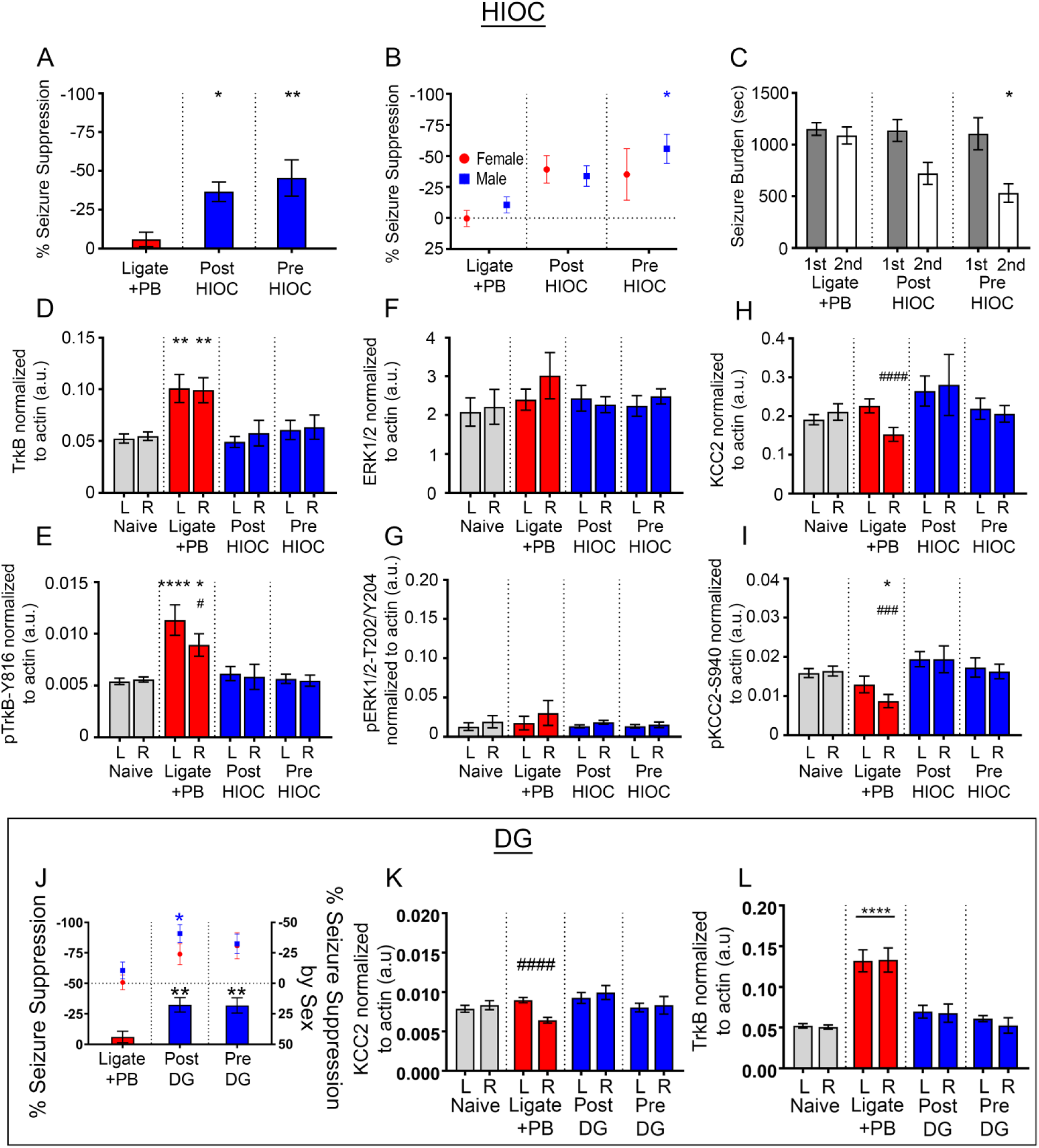
HIOC and DG significantly rescued PB-refractory seizures at P7. (A) EEG Percent seizure suppression for Ligate+PB, Post HIOC, and Pre HIOC treated P7 pups. * p<0.05, ** p<0.01 by one-way ANOVA. **(B)** EEG Percent seizure suppression by sex for Ligate+PB, Post HIOC, and Pre HIOC treated P7 pups. * p<0.05 by two-way ANOVA. **(C)** EEG seizure burdens for Ligate+PB, Post HIOC, and Pre HIOC treated P7 pups. * p<0.05 by two-way ANOVA. **(D)** Contralateral (L) and ipsilateral (R) TrkB expression 24h after ischemic insult for all treatment groups. ** p<0.01 by one-way ANOVA. All proteins were normalized to housekeeping protein β-actin. **(E)** Contralateral (L) and ipsilateral (R) pTrkB-Y816 expression 24h after ischemic insult for all treatment groups. * p<0.05. **** p<0.0001 by one-way ANOVA. # indicated differences between contralateral and ipsilateral hemispheres within groups; # p<0.05 by two-tailed *t* test. **(F)** Contralateral (L) and ipsilateral (R) ERK1/2 expression 24h after ischemic insult for all treatment groups. **(G)** Contralateral (L) and ipsilateral (R) pERK1/2-T202/Y204 expression 24h after ischemic insult for all treatment groups. **(H)** Contralateral (L) and ipsilateral (R) KCC2 expression 24h after ischemic insult for all treatment groups. #### p<0.0001 by two-tailed *t* test. **(I)** Contralateral (L) and ipsilateral (R) pKCC2-S940 expression 24h after ischemic insult for all treatment groups. * p<0.05 by one-way ANOVA. ### p<0.001 by two-tailed *t* test. **(J)** EEG percent seizure suppression for Ligate+PB, Post DG, and Pre DG treatment groups plotted against left y-axis. ** p<0.01 by one-way ANOVA. EEG percent seizure suppression by sex for Ligate+PB, Post DG, and Pre DG treatment groups plotted against right y-axis. * p<0.05 by two-way ANOVA. Horizontal dotted line represents 0% seizure suppression on right y-axis. **(K)** Contralateral (L) and ipsilateral (R) KCC2 expression 24h after ischemic insult for all treatment groups. # signified hemispheric differences within treatment groups. #### p<0.0001 by two-tailed *t* test. **(L)** Contralateral (L) and ipsilateral (R) TrkB expression 24h after ischemic insult for all treatment groups. **** p<0.0001 by one-way ANOVA.

To evaluate the role of HIOC effects on TrkB-pathway activation, expression levels of TrkB, ERK1/2, and KCC2 were examined 24h post-ligation. Similar to LM intervention, both Post and Pre HIOC groups significantly rescued TrkB-pathway activation (Fig. 5D) and pTrkB-Y816 activation bilaterally (Fig. 5E). Total ERK1/2 expression was not significantly impacted by ischemic insult or any HIOC treatment (Fig. 5F) similar to LM intervention (Fig. 4H). In contrast to Pre LM intervention (Fig. 4I), Post HIOC and Pre HIOC groups did not demonstrate an increase in pERK1/2-T202/Y204 expression (Fig. 5G). In summary, the differences in LM vs. HIOC-mediated downstream ERK1/2 phosphorylation may depend upon site-specific TrkB binding.

KCC2 expression decreased unilaterally in the ipsilateral ischemic hemisphere in the Ligate+PB group (Fig. 5H). Both Post HIOC and Pre HIOC groups demonstrated significant rescue of KCC2 and pKCC2-S940 (Fig. 5I) expression in the ipsilateral hemisphere. In summary, HIOC and LM had similar effects on TrkB, pTrkB-Y816, KCC2, pKCC2-S940, and ERK1/2 expression. In contrast to LM, HIOC treatment did not result in activation of the pERK1/2-T202/Y204 downstream pathway.

Treatment of P7 ischemic pups with another full TrkB agonist, DG, also significantly rescued PB-refractoriness in the Post DG and Pre DG groups (Fig. 5J). Male pups in the Post DG group had significantly greater seizure suppression than male pups in the Ligate+PB group (Fig. 5J), in contrast to both LM and HIOC treatments. No significant sex differences in seizure suppression were noted for both full TrkB agonists HIOC and DG. Comparing both of these TrkB agonists to LM highlights that male and female pups differentially responded to BDNF loop II mimetics and full TrkB agonists. However, the post-ischemic KCC2 degradation and TrkB pathway activation was rescued in the Post and Pre DG groups (Fig. 5K, L) similar to interventions with LM and HIOC. The differential activation of the TrkB-ERK1/2 pathway downstream of BDNF-TrkB signaling elicited by the BDNF mimetic LM was not detected with the TrkB agonists HIOC and DG.

### Differences of developmental profiles of TrkB and PLCγ1 expression in naïve CD-1 pups by sex

At P5, females expressed significantly higher levels of TrkB in the cortex compared to males (Fig. 6A1, A2 and B). Developmentally, TrkB expression in the cortex of female pups declined significantly from P5 to P21. In contrast, TrkB expression in males did not decline significantly from P5 to P21. Both female and male pups showed significant decline of pTrkB-Y816 expression from P5 to P21 (Fig. 6C). In the cortex, ratios of pTrkB-Y816 to total TrkB were not significantly different by sex or age (Fig. 6D).

**Fig. 6.**
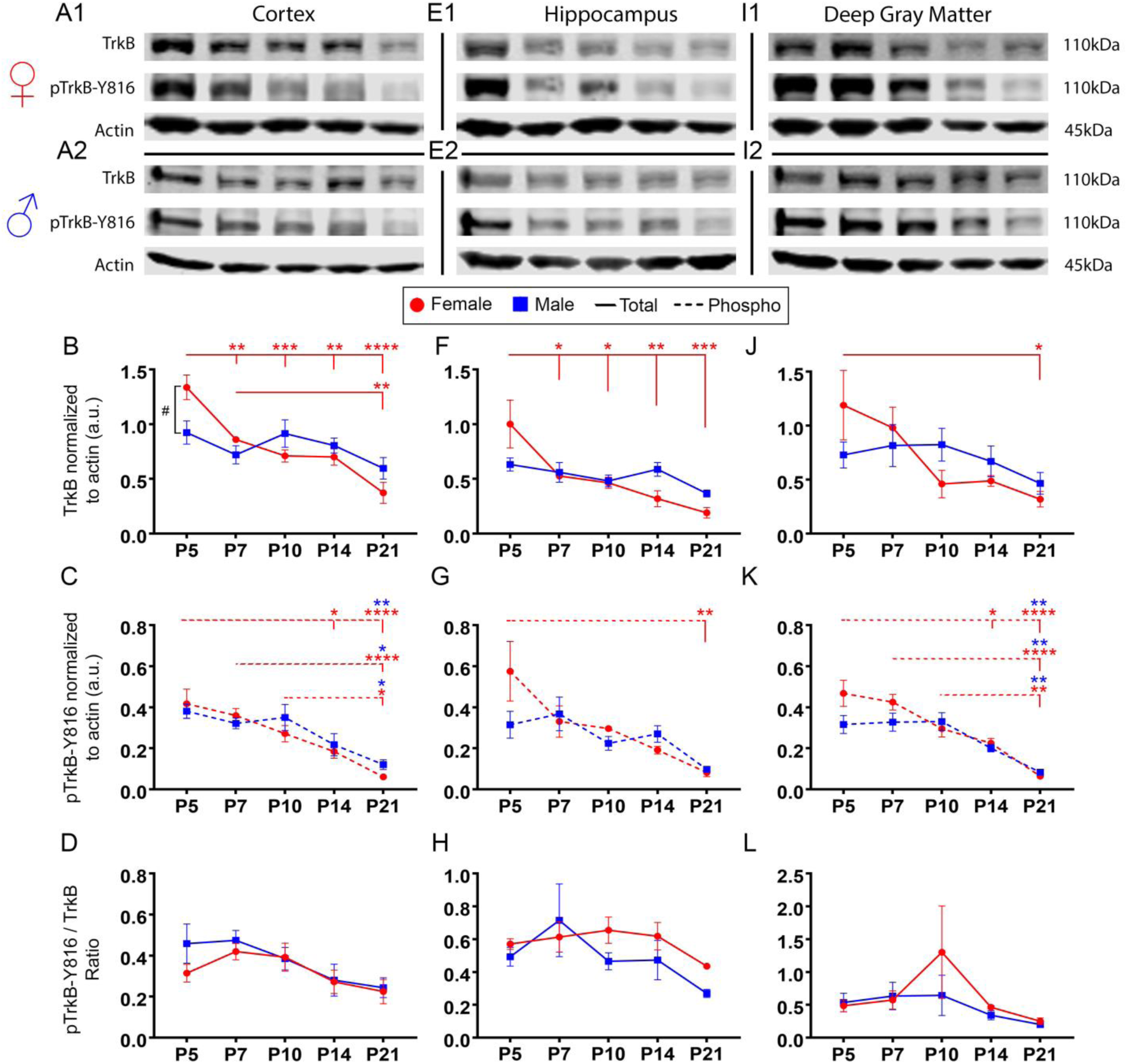
TrkB expression significantly decreased from P5 to P21 in a region-specific manner in naïve female pups. All proteins were normalized to β-actin. **(A1)** Representative Western blots showing TrkB and pTrkB-Y816 expression in cortex for all female and (**A2**) male age groups. **(B)** TrkB expression in cortical tissue from P5 to P21. ** p<0.01, *** p<0.001, **** p<0.0001 by two-way ANOVA. **(C)** pTrkB-Y816 expression in cortical tissue from P5 to P21. # p<0.05 signified difference between sexes at a given age. * p<0.05, ** p<0.01, **** p<0.0001 by two-way ANOVA. **(D)** pTrkB-Y816 normalized to total TrkB in cortical tissue from P5 to P21. **(E1)** Representative Western blots showing TrkB and pTrkB-Y816 expression in hippocampus for all female and (**E2**) male age groups. **(F)** TrkB expression in hippocampal tissue from P5 to P21. * p<0.05, ** p<0.01, *** p<0.001 by two-way ANOVA. **(G)** pTrkB-Y816 expression in hippocampal tissue from P5 to P21. ** p<0.01 by two-way ANOVA. **(H)** pTrkB-Y816 normalized to total TrkB in cortical tissue from P5 to P21. **(I1)** Representative Western blots showing TrkB and pTrkB-Y816 expression in deep gray matter for all female and (**I2**) male age groups. **(J)** TrkB expression in deep gray matter from P5 to P21. * p<0.05 by two-way ANOVA. **(K)** pTrkB-Y816 expression in deep gray matter from P5 to P21. * p<0.05, ** p<0.01, **** p<0.0001 by two-way ANOVA. **(L)** pTrkB-Y816 normalized to total TrkB in deep gray matter from P5 to P21.

**Fig. 7.**
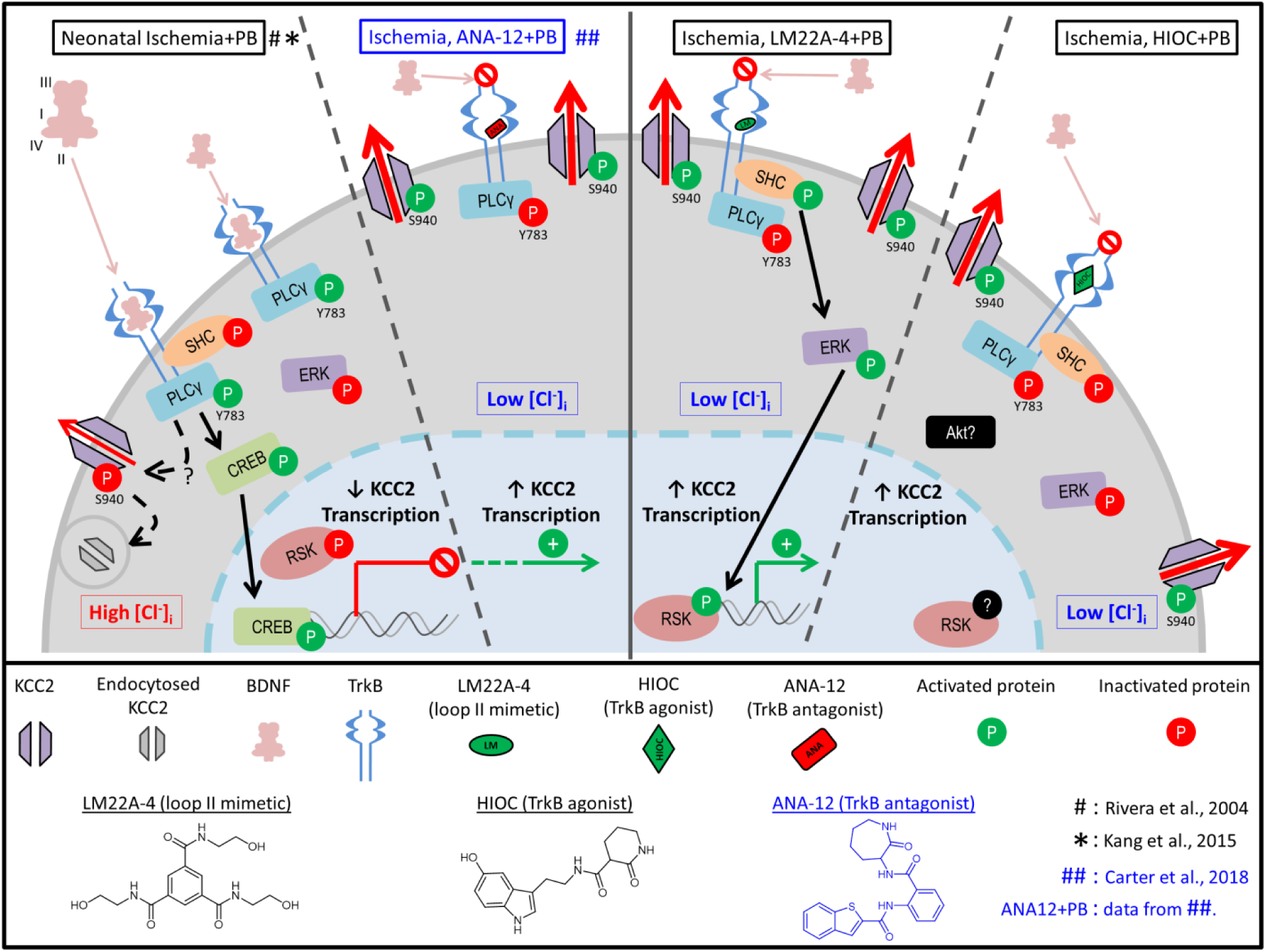
Summary schematic of TrkB signaling pathways following neonatal ischemia. Post-ischemia endogenous BDNF release results in activation of the TrkB-PLCγ1 pathway, thereby down-regulating KCC2 expression (data summarized from *13*, *40*) 24h post-ischemia. Treatment with the small-molecule TrkB antagonist ANA12 rescued post-ischemic TrkB-PLCγ1 pathway activation-mediated KCC2 degradation (data summarized from (*5*)). Intervention with LM22A-4 a TrkB partial agonist also rescued TrkB-PLCγ1 pathway activation similar to ANA12, and activated the TrkB-ERK1/2 pathway instead. Treatment with full TrkB agonist HIOC replicated the LM findings and rescued TrkB-PLCγ1 pathway activation but did not activate the TrkB-ERK1/2 pathway indicating TrkB site-specific engagement dictate downstream cascades.

Similar to the results found in the cortex, TrkB expression in the hippocampi of female pups declined significantly from P5 to P21, whereas TrkB expression in males did not decline significantly (Fig. 6E1, E2, and F). pTrkB-Y816 expression also declined significantly in the hippocampi of female pups from P5 to P21 (Fig. 6G). Ratios of pTrkB-Y816 to total TrkB were not sexually dimorphic and did not change significantly between ages of P5 to P21 in the hippocampus (Fig. 6H). In deep gray matter, TrkB expression decreased significantly between P5 and P21 in females, whereas TrkB expression remained stable and did not decline significantly in males (Fig. 6I1, I2, and J). pTrkB-Y816 expression decreased significantly in both females and males from P5 to P21 (Fig. 6I1, I2, and K). However, ratios of pTrkB-Y816 to total TrkB were not different between sexes sexually dimorphic and did not change significantly between ages of P5 to P21 in the deep gray matter (Fig. 6L). These results suggest that sexually dimorphic expression levels of TrkB in the cortex may underlie the sexually dimorphic seizure susceptibility and rescue of PB-refractoriness with LM (Fig. 1 and 2), as neonatal seizures are cortical. Furthermore, the attenuation of pTrkB-Y816 from P5 to P21 in both males and females suggests that TrkB pathway activation decreases with age.

At P5, females showed significantly higher PLCγ1 and pPLCγ1-Y783 expression in the cortex than at P21 (Fig. S5A and B). In contrast, PLCγ1 and pPLCγ1-Y783 expression in males was not significantly different between age groups. The hippocampus and deep gray matter did not show significant differences between sexes in PLCγ1 or pPLCγ1-Y783 expression during development (Fig. S5D, E, G, and H). Similarly, ratios of pPLCγ1-Y783 to total PLCγ1 were not significantly different between sexes during development for all examined brain regions (Fig. S5C, F, and I).

### Differences in developmental expression profile of KCC2 in naïve CD-1 pups by sex

Males showed a significant increase in KCC2 expression between P5 and P21, and P7 and P21, whereas females did not (Fig. S5J and K). pKCC2-S940 was not found to be significantly dephosphorylated in both sexes (Fig. S5J and K). In contrast, only females showed significantly lower ratios of pKCC2-S940 to total KCC2 in the cortex (Fig. S5L). In the hippocampus, both males and females did not show a significant change in KCC2 expression (Fig. S5M and N). pKCC2-S940 expression and ratios of pKCC2-S940 to total KCC2 were not found to be sexually dimorphic between ages (Fig. S5M-O). In deep gray matter, both females and males showed significantly lower expression of pKCC2-S940 at P21 (Fig. S5P-R). In summary, the developmental age-dependent decline in KCC2 expression demonstrated an opposite trend to TrkB expression during the same developmental period.

## Discussion

The main findings of this study are: 1. A single-dose of LM, a small-molecule BDNF loop II mimetic, significantly rescued PB-refractoriness in a mouse model of neonatal seizures. 2. LM was more efficacious in female pups at P7. 3. LM prevented ischemia-induced TrkB-PLCγ1 pathway activation and subsequent KCC2 degradation while significantly increasing ERK1/2 phosphorylation. 4. The full TrkB agonists HIOC and DG also rescued PB-refractoriness and ischemia-induced KCC2 degradation, indicating that the efficacy of LM was not site-dependent. HIOC also prevented TrkB-PLCγ1 pathway activation and KCC2 degradation, however without increasing ERK1/2 phosphorylation. 5. At P10, seizures responded efficaciously to PB, indicating age-dependent emergence of refractory neonatal seizures (P7 vs. P10) and were not associated with significant BDNF-TrkB pathway activation. 6. Chemogenetic inactivation of TrkB receptor in P7 pups resulted in a significant reduction in their post-ischemic seizure susceptibilities supporting the role of the BDNF-TrkB pathway activation in aggravation of neonatal seizures. 7. The developmental expression profiles demonstrated a significant decline in TrkB and PLCγ1 expression driven by female pups, versus a significant increase in KCC2 expression driven by male pups from P5 to P21. 8. Early in development, females showed significantly higher TrkB expression in the cortex, which may underlie the differences in LM anti-seizure efficacy by sex.

### LM, a BDNF loop II mimetic, functioned as a TrkB antagonist for endogenous BDNF following ischemia

The therapeutic potential of neurotrophin-based treatments for neurological diseases is an active field of preclinical research (*38*). In contrast to the role of neurotrophins in adulthood, the role of neurotrophin-based interventions in the developing neonatal brain is only recently emerging (*39, 40*). BDNF-TrkB signaling is age-dependent, as BDNF robustly activates TrkB in the neonatal rodent brain but is this interaction is attenuated in the adult rodent brain (*41*). In human frontal cortex, BDNF expression may gradually decrease with age (*42*). Following BDNF binding, TrkB forms a homodimer and auto-phosphorylates Tyr residues in its intracellular domain (*43*). The phosphorylation of the TrkB homodimer residues initiates multiple downstream signal transduction pathways that affect neuronal survival, synaptogenesis, dendritic structure, and activity-dependent synaptic plasticity in a cell-type specific manner (*44–46*). Further, these signaling pathways are dependent upon the time course of BDNF-TrkB activation, as acute vs. chronic activation of BDNF-TrkB are associated with divergent outcomes (*45, 47, 48*). The complexity of BDNF function is apparent in its role as a selective regulator of gene expression via its modulation of RNA-binding proteins, and micro-RNAs [reviewed in (*49*)]. Additionally, both the human and rodent BDNF gene consist of 9 exons, each with their own promoters resulting in at least 10 different transcripts [reviewed in (*50*)]. These alternative *Bdnf* transcripts undergo unique temporal and spatial modulation that allow different factors, such as hypoxic response elements, to regulate BDNF signaling in a cell-type and circuit-specific manner (*39, 51*). The overexpression of a cleavage resistant precursor of BDNF (proBDNF) has been demonstrated to reduce KCC2 protein expression via the p75 neurotrophin receptor (*52*), further highlighting the complexity of BDNF-TrkB signaling in health and disease.

Downstream activation of the TrkB-PLCγ1-pathway has been tied to KCC2 hypofunction by several studies (*11, 12, 53*). In an adult model of limbic epilepsy, prevention of PLCγ1-pathway activation prevented epileptogenesis (*54, 55*). Further, AAV-*Cre* mediated reduction of KCC2 in the CA1 and dentate gyrus of the adult mouse hippocampus resulted in some of the core phenotypes of medial temporal lobe epilepsy, such as spontaneous seizures, gliosis, and neuronal loss (*56*). These results from adult mouse models of epilepsy, and the prevalence of pathogenic KCC2 mutations in human epilepsy (*57*) support the critical role of KCC2 and pathways that promote its hypofunction in epilepsy.

Previous work in the neonatal brain has demonstrated the importance of preventing BNDF-TrkB activation following excitotoxic injury when using a small-molecule TrkB antagonist ANA12 [(*5*), see schematic Fig 7]. ANA12 prevented the activation of the TrkB-PLCγ1 pathway, reversed post-ischemic KCC2 hypofunction, and rescued P7 PB-refractory seizures (*2, 5*). In this same neonatal mouse model, the BDNF loop II mimetic LM also prevented BDNF-mediated TrkB-PLCγ1 pathway activation and KCC2 hypofunction. In conjunction with the proposed binding of LM to TrkB (*14*), and the observation that LM intervention was efficacious, our data suggest that the presence and subsequent binding of LM to TrkB also prevents the cascade of endogenous post-ischemic BDNF-TrkB signaling similar to the TrkB antagonist ANA12. Taken together these data indicate that the BDNF loop II mimetic LM and TrkB antagonist ANA12 both act as TrkB antagonists to the endogenous BDNF released in the post-ischemic neonatal brain (*2, 5, 15*).

### TrkB-ERK1/2 pathway modulation by the BDNF loop II mimetic

LM binding to TrkB lead to the activation of the TrkB-ERK1/2 pathway, similar to previous reports from in vitro studies that peptide mimetics of either loop I, III, or IV of BDNF can induce AKT and ERK phosphorylation (*58, 59*). TrkB activation also phosphorylates residue Y816 on its intracellular domain, which permits recruitment and activation of the PLCγ1 pathway, which activates numerous pathways including the Ca^2+^/calmodulin-dependent kinases (*60, 61*). In this study, LM rescued the ischemia-induced phosphorylation of residue Y816 on TrkB, suggesting that the prevention of the TrkB-PLCγ1 pathway mediated KCC2 degradation occurred by LM binding to TrkB.

It has been shown that the use of HIOC in vivo protected retinas from excitotoxic retinal degeneration in a TrkB-dependent manner (*16*). Although both LM and HIOC significantly rescued PB-refractoriness at P7, HIOC functioned differently than LM as it did not induce activation of the TrkB-ERK1/2 pathway. These results indicate that TrkB agonists can prevent emergence of refractory seizures by preventing the pathological BDNF-TrkB-PLCγ1 pathway activation in post-ischemic neonatal brains, thus functionally acting as antagonists.

The ERK1/2 and AKT pathways are known to promote neurogenesis through the Ras signaling cascade (*61*), which regulates a multitude of processes, including cell migration, differentiation, proliferation, and transcription (*62, 63*). Previous literature has demonstrated that LM promoted cell survival in a model of nonarteritic anterior ischemic optic neuropathy (*64*). Supporting these results, the Pre LM group showed a significant increase in pERK1/2-T202/Y204 expression. Intervention with BDNF loop II mimetic LM was shown to have beneficial long-term effects in vivo, particularly in rescuing adult post-traumatic cortical epileptogenesis in a TrkB-dependent manner (*65*), supporting the findings reported in this study.

#### Evidence for BDNF hyperactivity in neurological disorders

In autism, a severe neurodevelopmental disorder with pathogenesis that occurs during the neonatal period, the early hyperactivity of BDNF may play an important role as high BDNF levels have been reported in neonatal blood samples from children with autistic spectrum disorders (*66, 67*). Valproic acid (VPA) exposure during pregnancy increases the risk of congenital malformations and autism (*68–70*). VPA administration to pregnant rodents during the 2^nd^ week of gestation is a well-investigated model of autism (*71*). The VPA model of autism has demonstrated increased BDNF (*71*), increased neuronal intracellular chloride concentration (*72*), and a disruption of the GABA developmental sequence (*72, 73*). Research utilizing a preclinical model of Fragile X syndrome, an inherited form of intellectual disability and autism spectrum disorder, administered LM to neonatal *Fmr1* KO mice and rescued cell type specific cortical developmental alterations (*74*). These studies suggest that TrkB agonists have a potential role in neurodevelopmental disorders with high levels of BDNF such as autism.

TrkB agonists can differentially activate selective downstream pathways, and TrkB-PLCγ1 activation is complex. This concept is already evident in recent literature that has shown evidence of TrkB receptor activation and signaling without dimerization, suggesting that TrkB can under certain conditions exist and function as a monomeric receptor at the plasma membrane (*75*). Moreover, recent studies examining plasma membrane diffusion kinetics have demonstrated that up to 20% of the Trk family may exist as dimers or oligomers prior to neurotrophin binding (*76*), though their function remains a topic of debate (*77*). These observations warrant further investigation of the complexities involved in neurotrophin-based signaling in vivo. One specific example of the in vivo complexity of BDNF-TrkB signaling is in the superoxide dismutase 1 mouse model of familial ALS (*78*). In this model, the complete deletion of TrkB in motor neurons in vivo slowed the progression of the disease and permitted the maintenance of motor function (*78*). Another indication of the complexity of BDNF-TrkB signaling is in vitro studies that have identified that an increase in BDNF renders motor neurons more susceptible to excitotoxic insults in a TrkB dependent manner, however, not all motor neurons were protected by blocking TrkB (*79*).

### Sexually dimorphic developmental profiles of TrkB and KCC2

TrkB signaling is known to be sexually dimorphic in a region-specific manner (*80–82*). In BDNF^+/-^ mice, greater levels of pTrkB-Y705 activation were shown in the frontal cortex and striatum of male mice 10-16 weeks of age compared to females. These changes in TrkB phosphorylation state were accompanied by concomitant increases in ERK1/2 phosphorylation(*80*), suggesting a differential signaling mechanism that was sexually dimorphic. The developmental expression of KCC2 is also known to increase with age, and this developmental profile is sex-dependent, with female pups expressing higher levels than male pups (*15, 83, 84*). Similarly, rodent models have demonstrated higher female expression of KCC2 mRNA compared to age-matched males in the substantia nigra and mediobasal hypothalamus (*83, 85*).

## Conclusion

The findings reported here demonstrate that the BDNF loop II mimetic LM and full TrkB agonists HIOC and DG, significantly rescued PB-refractoriness and prevented post-ischemic degradation of KCC2. Sex-dependent differences in developmental profiles for TrkB and KCC2 were identified that may underlie the significant sex-dependent variance in efficacy noted for LM and the full TrkB agonists. Additionally, the TrkB receptor plays a unique role in post-ischemic seizure susceptibility in the neonatal brain as shown using chemogenetic techniques. In summary, results of this study and previous results from a small-molecule TrkB antagonist ANA12 (*2, 5*) indicate that refractory seizures following neonatal ischemia can be rescued acutely by both by TrkB antagonists and agonists. These findings indicate that under neonatal ischemic conditions, the TrkB agonists investigated here pharmacologically acted similar to TrkB antagonists by preventing cascades associated with the endogenous BDNF-TrkB pathway activation, providing novel insights into post-ischemic pathophysiology in immature brains. This highlights the crucial role of the TrkB receptor in neonatal seizure susceptibility and emergence of refractory seizures.

## Supporting information

Graphical abstract

## Acknowledgments

### Funding

This work was supported by the Eunice Kennedy Shriver National Institute of Child Health and Human Development of the National Institutes of Health under Award Number R01HD090884 (SDK). The content is solely the responsibility of the authors and does not necessarily represent the official views of the National Institutes of Health.

### Author contributions

SDK designed research; PAK, BJS, BMC, and SDK performed research; PAK, BJS, and SDK analyzed data; PAK, BJS, and SDK wrote the paper.

## Supplementary Materials

**Fig. S1.**
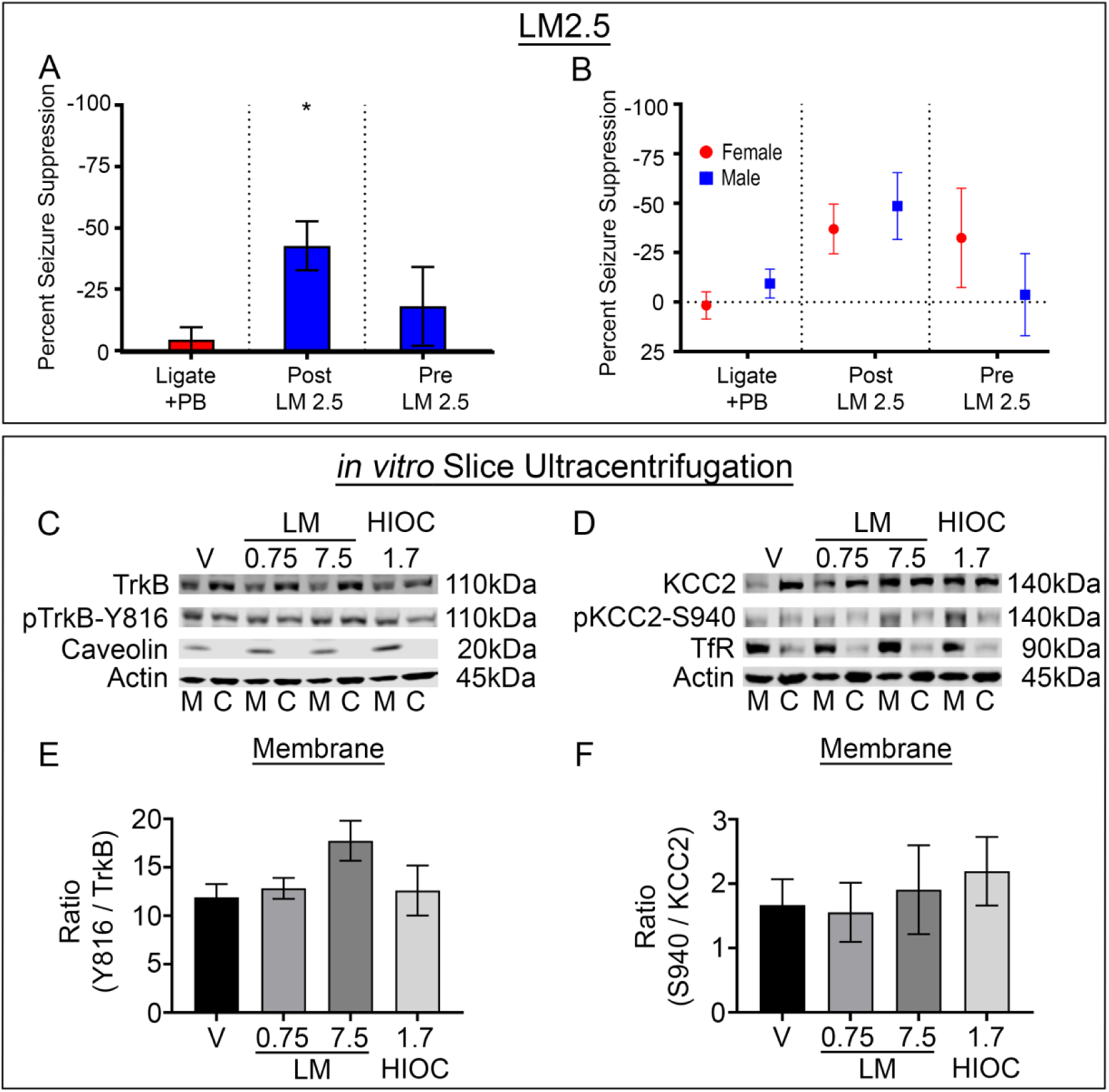
LM graded dose failed to significantly improve on in vivo anti-seizure efficacy and in vitro TrkB phosphorylation in naïve P7 brain slices. **(A)** EEG percent seizure suppression for Ligate+PB, Post LM 2.5, and Pre LM 2.5 treated P7 pups. * p<0.05 (Post LM2.5 vs. Ligate+PB) by one-way ANOVA. **(B)** EEG percent seizure suppression by sex for Ligate+PB, Post LM 2.5, and Pre LM 2.5 treated P7 pups. **(C)** Representative Western blot showing membrane and cytosolic TrkB and pTrkB-Y816 expression after incubation with TrkB agonists. **(D)** Representative Western blot showing membrane and cytosolic KCC2 and pKCC2-S940 expression after incubation with TrkB agonists. **(E)** Ratio of pTrkB-Y816 to total TrkB at the plasma membrane. **(F)** Ratio of pKCC2-S940 to total KCC2 at the plasma membrane.

**Fig. S2.**
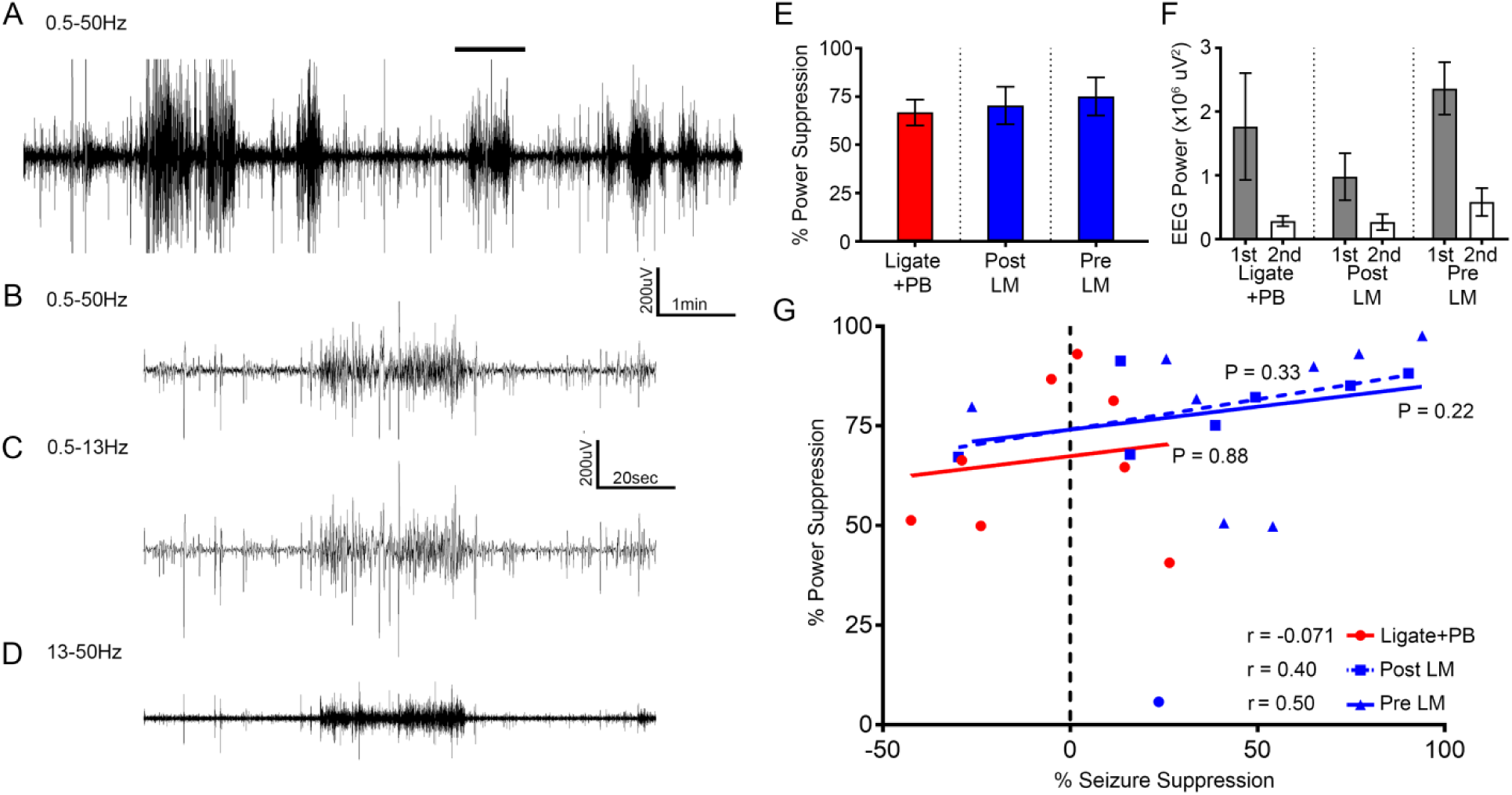
EEG power alone failed to identify the LM-mediated rescue of PB-refractoriness in P7 pups. **(A)** Representative raw 10 min EEG trace from 0.5-50Hz of refractory ischemic seizures from a neonatal P7 mouse pup. **(B)** A single ictal event from **A** (solid bar – 2 min expanded timescale raw trace). **(C and D)** Filtered EEG trace of the same ictal event in **B** filtered to show low frequency and high frequency components of the same ictal event (0.5-13Hz and 13-50Hz). **(E)** EEG percent power suppression for Ligate+PB, Post LM, and Pre LM treated P7 pups. **(F)** EEG 1^st^ and 2^nd^ hour EEG powers for Ligate+PB, Post LM, and Pre LM treated P7 pups. **(G)** Percent power suppression plotted as a function of percent seizure suppression. Post-hoc comparisons were performed using Spearman’s two-tailed nonparametric test.

**Fig. S3.**
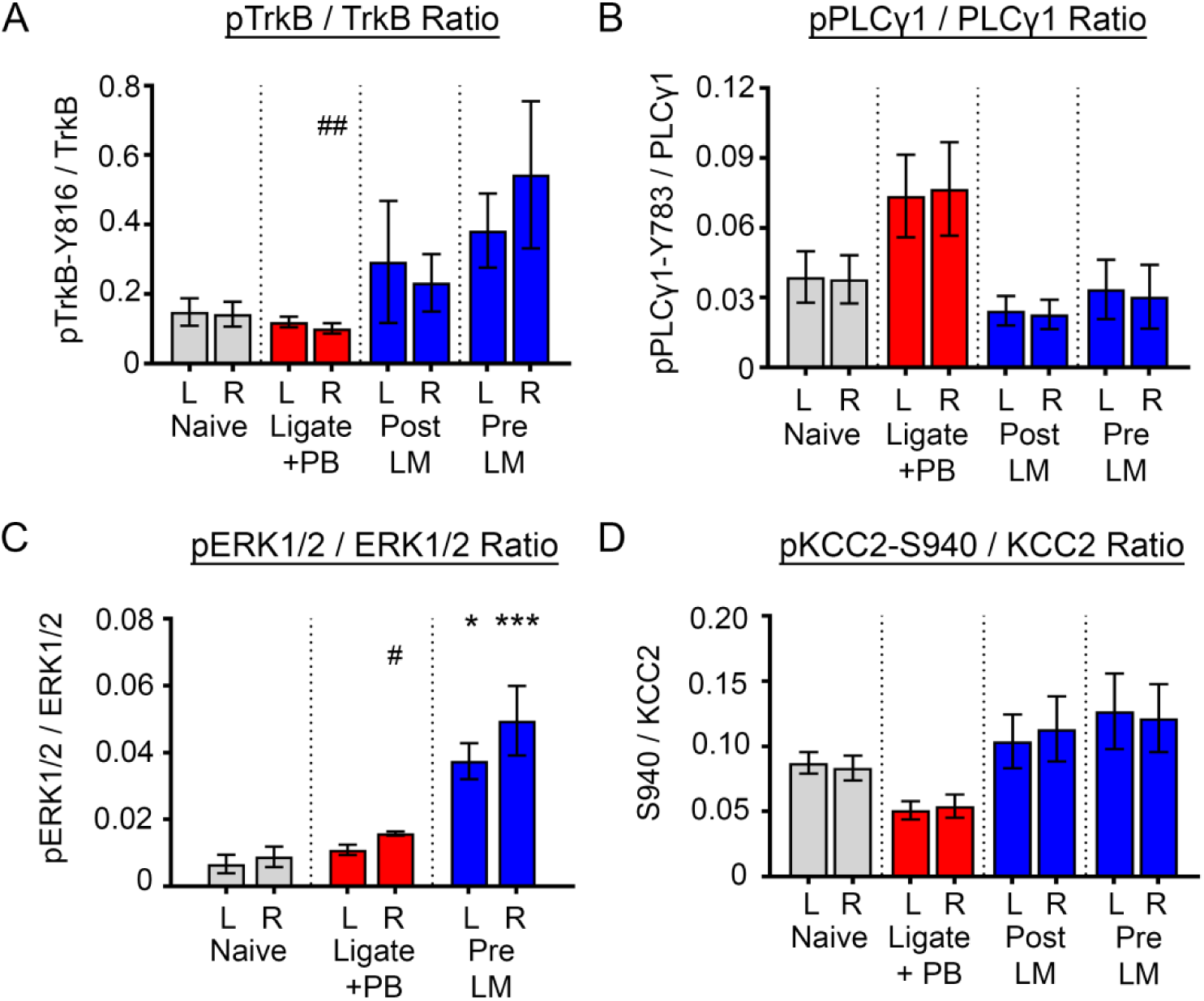
Normalization of phospho-proteins to their total proteins in P7 ischemic pups. **(A)** pTrkB-Y816 normalized to total TrkB for Naïve, Ligate+PB, Post LM, and Pre LM at P7. # signified hemispheric differences within groups, ## p<0.01 by two-tailed *t* test. **(B)** pPLCγ1-Y783 normalized to total PLCγ1 for Naïve, Ligate+PB, Post LM, and Pre LM at P7. **(C)** pERK1/2 normalized to total ERK1/2 for Naïve, Ligate+PB, and Pre LM at P7. * p<0.05, *** p<0.001 by one-way ANOVA. # p<0.05 by two-tailed *t* test. **(D)** pKCC2-S940 normalized to total KCC2 for Naïve, Ligate+PB, Post LM, and Pre LM at P7.

**Fig. S4.**
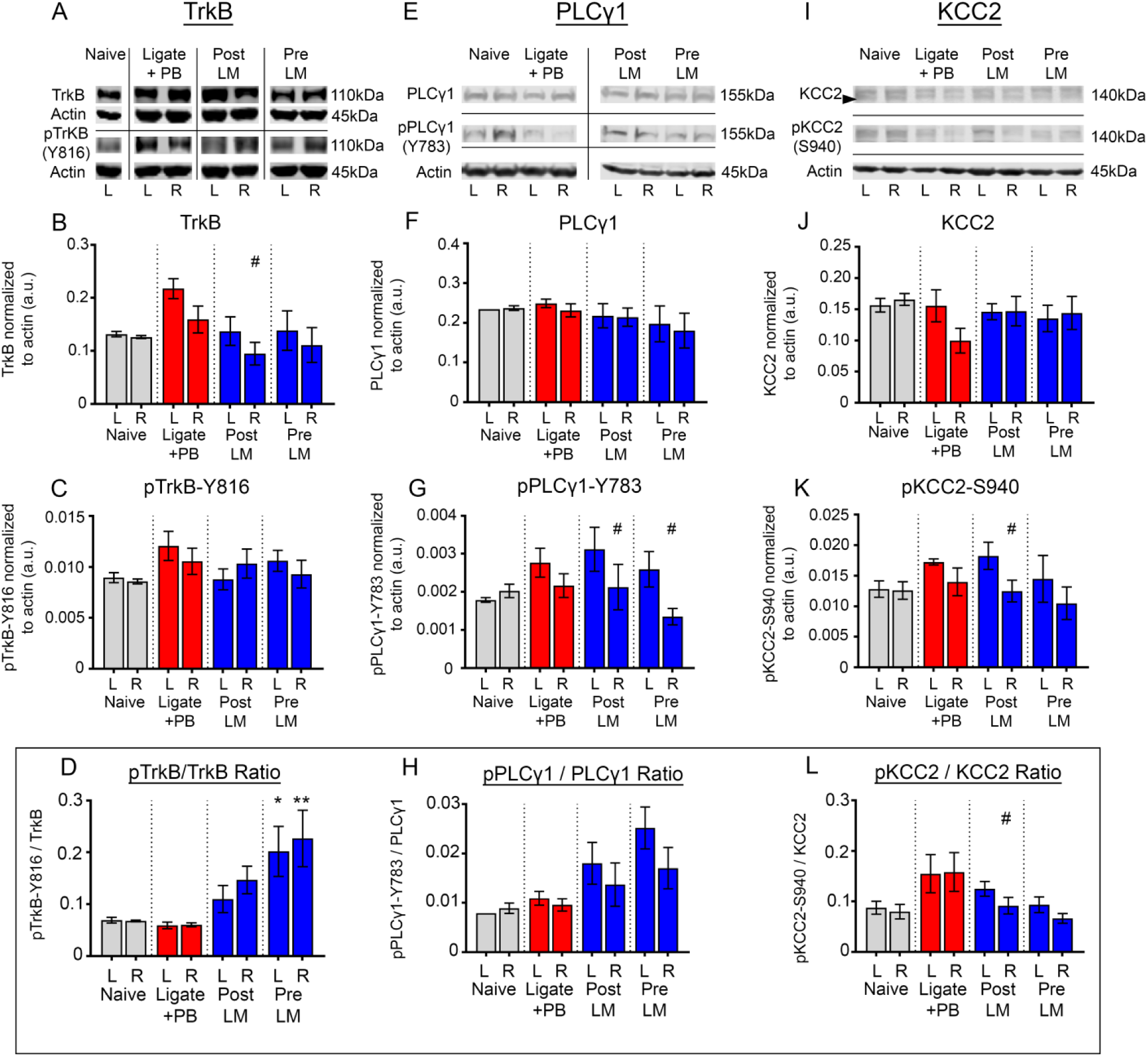
TrkB-pathway activation was not significant in P10 ischemic pups. All proteins of interest were normalized to housekeeping protein β-actin. **(A)** Representative Western blots showing TrkB and pTrkB-Y816 expression in P10 pups. **(B)** Contralateral (L) and ipsilateral (R) TrkB expression 24h after ischemic insult for all treatment groups. # signified hemispheric differences within groups. # p<0.05 by two-tailed *t* test. **(C)** Contralateral (L) and ipsilateral (R) pTrkB-Y816 expression 24h after ischemic insult for all treatment groups. **(D)** pTrkB-Y816 normalized to total TrkB for Naïve, Ligate+PB, Post LM, and Pre LM at P10. * p<0.05, ** p<0.01 by one-way ANOVA. **(E)** Representative Western blots showing PLCγ1 and pPLCγ1-Y783 expression. **(F)** Contralateral (L) and ipsilateral (R) PLCγ1 expression 24h after ischemic insult for all treatment groups. **(G)** Contralateral (L) and ipsilateral (R) pPLCγ1-Y783 expression 24h after ischemic insult for all treatment groups. # signified hemispheric differences within groups. # p<0.05 by two-tailed *t* test. **(H)** pPLCγ1-Y783 normalized to total PLCγ1 for Naïve, Ligate+PB, Post LM, and Pre LM at P10. **(I)** Representative Western blots showing KCC2 and pKCC2-S940 expression. **(J)** Contralateral (L) and ipsilateral (R) KCC2 expression 24h after ischemic insult for all treatment groups. **(K)** Contralateral (L) and ipsilateral (R) pKCC2-S940 expression 24h after ischemic insult for all treatment groups. # p<0.05 by two-tailed *t* test. **(L)** KCC2 normalized to total pKCC2-S940 for Naïve, Ligate+PB, Post LM, and Pre LM at P10. # signified hemispheric differences within groups. # p<0.05.

**Fig. S5.**
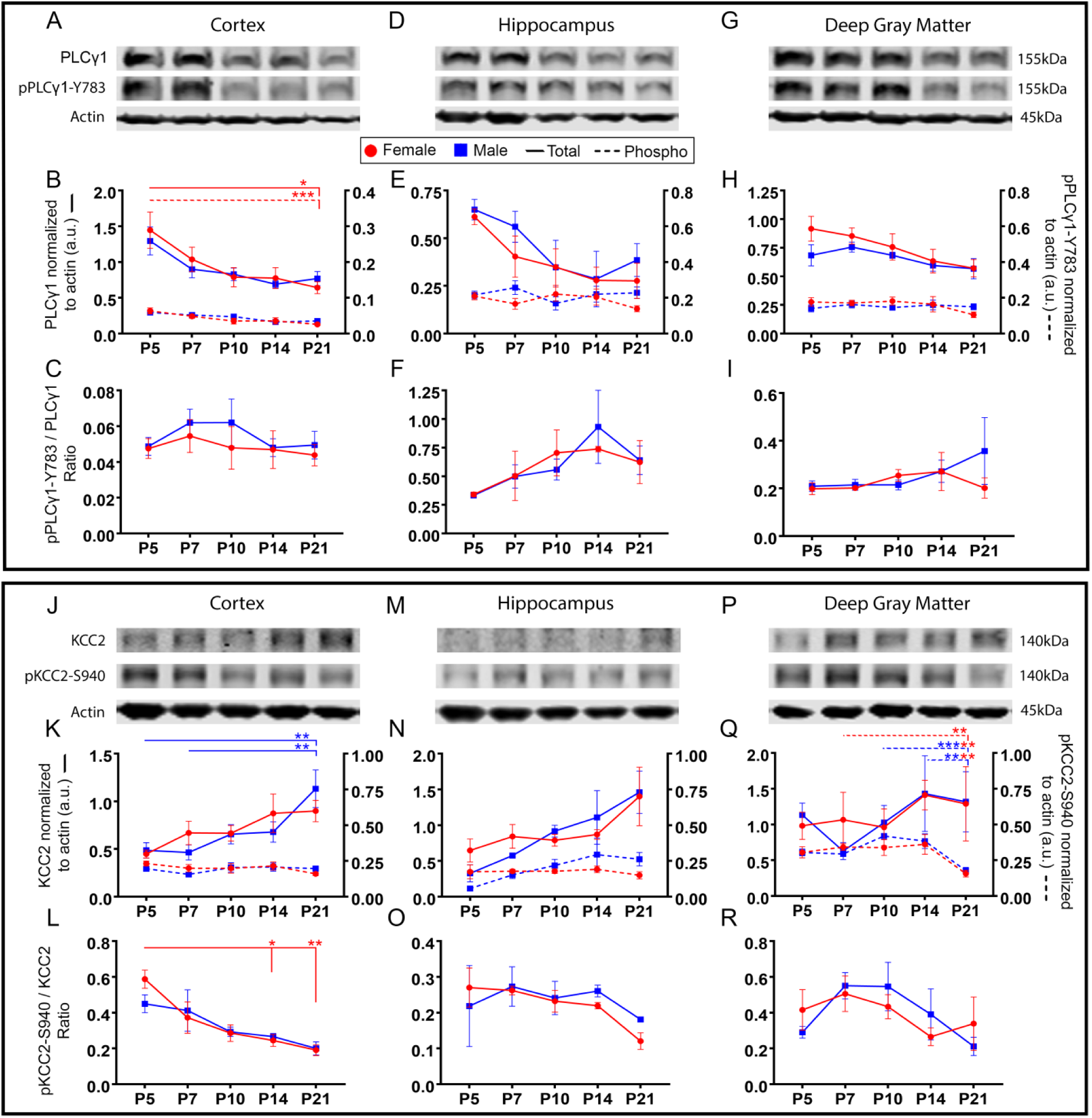
PLCγ1 and pPLCγ1-Y783 expression decreased significantly in female cortices, whereas KCC2 expression significantly increased in male cortices. All proteins were normalized to β-actin. **(A)** Representative Western blots showing PLCγ1 and pPLCγ1-Y783 expression in cortical tissue. **(B)** PLCγ1 and pPLCγ1-Y783 expression in cortical tissue from P5 to P21. * p<0.05, *** p<0.001, **** p<0.0001 by two-way ANOVA. **(C)** pPLCγ1-Y783 normalized to total PLCγ1 in cortical tissue from P5 to P21. **(D)** Representative Western blots showing PLCγ1 and pPLCγ1-Y783 expression in hippocampal tissue. **(E)** PLCγ1 and pPLCγ1-Y783 expression in hippocampal tissue from P5 to P21. **(F)** pPLCγ1-Y783 normalized to total PLCγ1 in hippocampal tissue from P5 to P21. **(G)** Representative Western blots showing PLCγ1 and pPLCγ1-Y783 expression in deep gray matter. **(H)** PLCγ1 and pPLCγ1-Y783 expression in deep gray matter from P5 to P21. **(I)** pPLCγ1-Y783 normalized to total PLCγ1 in hippocampal tissue from P5 to P21. **(J)** Representative Western blots showing KCC2 and pKCC2-S940 expression in cortical tissue. **(K)** KCC2 and pKCC2-S940 expression in cortical tissue from P5 to P21. ** p<0.01 by two-way ANOVA. **(L)** pKCC2-S940 normalized to total KCC2 in cortical tissue from P5 to P21. * p<0.05, ** p<0.01 by two-way ANOVA. **(M)** Representative Western blots showing KCC2 and pKCC2-S940 expression in hippocampal tissue. **(N)** KCC2 and pKCC2-S940 expression in cortical tissue from P5 to P21. **(O)** pKCC2-S940 normalized to total KCC2 in hippocampal tissue from P5 to P21. **(P)** Representative Western blots showing KCC2 and pKCC2-S940 expression in deep gray matter. **(Q)** KCC2 and pKCC2-S940 expression in deep gray matter from P5 to P21. ** p<0.01, *** p<0.001 by two-way ANOVA. **(R)** pKCC2-S940 normalized to total KCC2 in deep gray matter from P5 to P21.

**Table S1.** **Sample sizes for experimental paradigms**

| Sample size of pups in post-ligation EEG recordings |  |  |  |  |
| --- | --- | --- | --- | --- |
| Pups | Ligate+PB | Post LM 0.25 | Pre LM 0.25 |  |
| P7 total (M/F) [litters] | 28 (16/12) [9] | 26 (14/12) [6] | 27 (14/13) [9] |  |
| P10 total (M/F) [litters] | 11 (6/5) [3] | 13 (6/7) [3] | 11 (5/6) [3] |  |
| Pups | Ligate+PB | Post LM 2.5 | Pre LM 2.5 |  |
| P7 total (M/F) [litters] | 28 (16/12) [7] | 8 (4/4) [2] | 8 (4/4) [2] |  |
| Pups | Ligate+PB | Post HIOC | Pre HIOC |  |
| P7 total (M/F) [litters] | 28(16/12) [7] | 6 (3/3) [2] | 8 (4/4) [2] |  |
| Pups | Ligate+PB | Post DG | Pre DG |  |
| P7 total (M/F) [litters] | 28 (16/12) [7] | 16 (8/8) [2] | 15 (10/5) [2] |  |
| Sample size of pups that underwent Western Blot analysis after EEG recordings |  |  |  |  |
| Pups | Naïve | Ligate+PB | Post LM 0.25 | Pre LM 0.25 |
| P7 total (M/F) [litters] | 13 (6/7) [9] | 19 (10/9) [7] | 13 (5/8) [6] | 22 (11/11) [9] |
| P10 total (M/F) [litters] | 4 (2/2) [3] | 9 (5/4) [3] | 12 (6/6) [3] | 11 (5/6) [3] |
| Pups | Naïve | Ligate+PB | Post LM2.5 | Pre LM2.5 |
| P7 total (M/F) [litters] | 13 (6/7) [9] | 19 (10/9) [7] | 8 (4/4) [2] | 8 (4/4) [2] |
| Pups | Naïve | Ligate+PB | Post HIOC | Pre HIOC |
| P7 total (M/F) [litters] | 13 (6/7) [9] | 19 (10/9) [7] | 5 (3/2) [2] | 6 (4/2) [2] |
| Pups | Naïve | Ligate+PB | Post DG | Pre DG |
| P7 total (M/F) [litters] | 13 (6/7) [9] | 19 (10/9) [7] | 6 (4/2) [2] | 7 (4/3) [2] |
| Sample size of pups that underwent Western Blot analysis in Developmental Series |  |  |  |  |
| Pups | Cortex | Hippocampus | Deep Gray Matter |  |
| P5 total (M/F) [litters] | 4 (2/2) [3] | 3 (1/2) [3] | 4 (2/2) [3] |  |
| P7 total (M/F) [litters] | 4 (2/2) [3] | 6 (3/3) [3] | 6 (3/3) [3] |  |
| P10 total (M/F) [litters] | 4 (2/2) [3] | 4 (2/2) [3] | 4 (2/2) [3] |  |
| P14 total (M/F) [litters] | 4 (2/2) [3] | 4 (2/2) [3] | 4 (2/2) [3] |  |
| P21 total(M/F) [litters] | 4 (2/2) [3] | 4 (2/2) [3] | 4 (2/2) [3] |  |
| Sample size of pups that underwent in vitro drug incubation and Western Blot analysis |  |  |  |  |
| Pups | 0.75mM LM | 7.5mM LM | 1.7mM HIOC |  |
| P7 total (M/F) [litters] | 2 (1/1) [1] | 2 (1/1) [1] | 2 (1/1) [1] |  |
| Sample size of pups from TrkB<sup>F616A</sup> litters |  |  |  |  |
| Pups | WT<sup>-/-</sup> + 1NMPP1 |  | F616A<sup>+/+</sup> + 1NMPP1 |  |
| P7 total (M/F) [litters] | 18 (9/9) [5] |  | 10 (6/4) [5] |  |

**Table S2.**
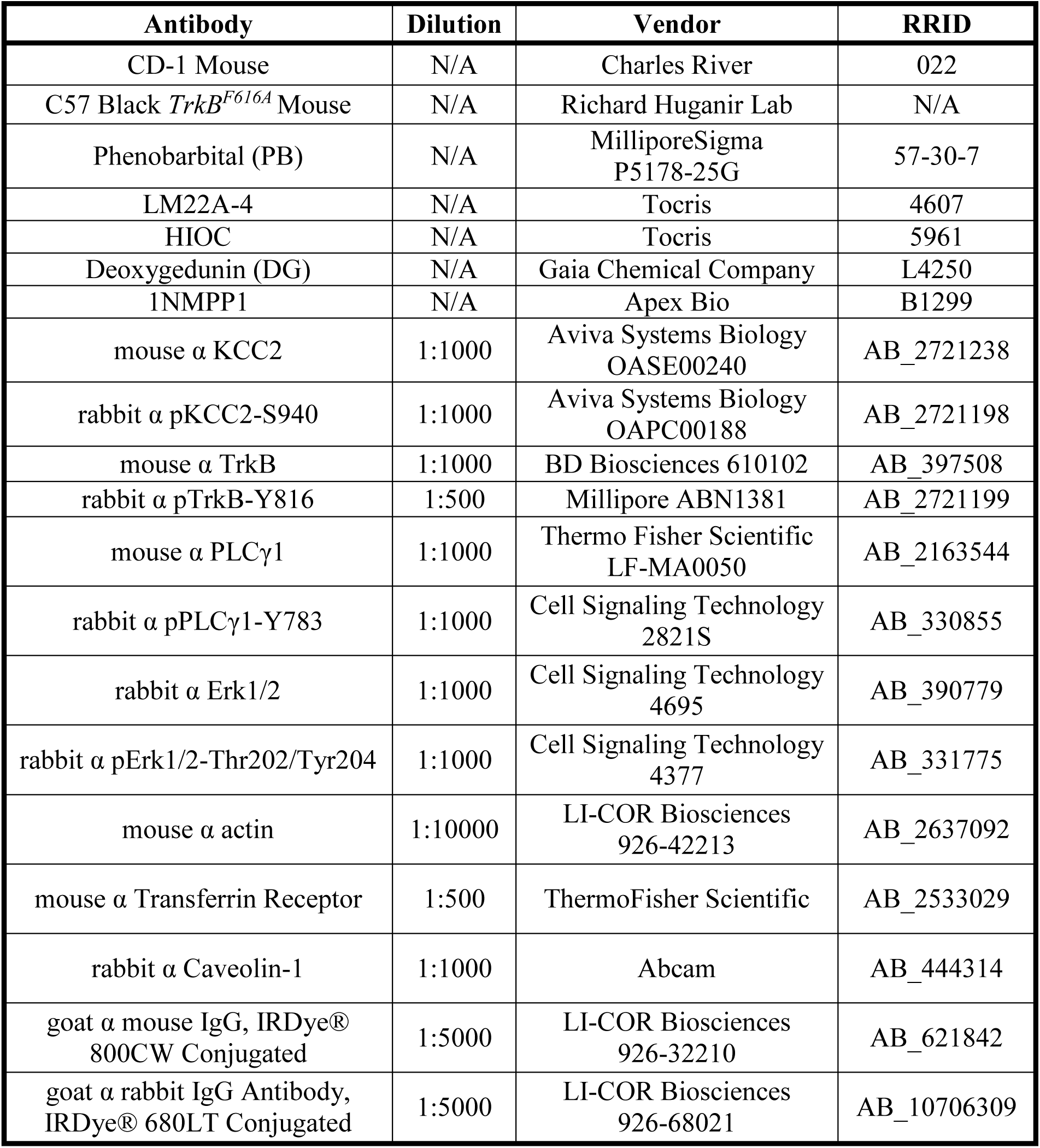
**Drugs, antibodies, and mice information**

## References and Notes

1. R. Aloyz, J. P. Fawcett, D. R. Kaplan, R. A. Murphy, F. D. Miller, Activity-Dependent Activation of TrkB Neurotrophin Receptors in the Adult CNS. Learn. Mem. 6, 216–231 (1999).

2. S. K. Kang, M. V. Johnston, S. D. Kadam, Acute TrkB inhibition rescues phenobarbital-resistant seizures in a mouse model of neonatal ischemia. Eur J Neurosci. 42, 2792–2804 (2015).

3. X.-P. He, L. Minichiello, R. Klein, J. O. McNamara, Immunohistochemical Evidence of Seizure-Induced Activation of trkB Receptors in the Mossy Fiber Pathway of Adult Mouse Hippocampus. J. Neurosci. 22, 7502–7508 (2002).

4. P. Kowiański, G. Lietzau, E. Czuba, M. Waśkow, A. Steliga, J. Moryś, BDNF: A Key Factor with Multipotent Impact on Brain Signaling and Synaptic Plasticity. Cell Mol Neurobiol. 38, 579–593 (2018).

5. B. M. Carter, B. J. Sullivan, J. R. Landers, S. D. Kadam, Dose-dependent reversal of KCC2 hypofunction and phenobarbital-resistant neonatal seizures by ANA12. Scientific Reports. 8, 11987 (2018).

6. H. H. C. Lee, J. A. Walker, J. R. Williams, R. J. Goodier, J. A. Payne, S. J. Moss, Direct Protein Kinase C-dependent Phosphorylation Regulates the Cell Surface Stability and Activity of the Potassium Chloride Cotransporter KCC2. J. Biol. Chem. 282, 29777–29784 (2007).

7. H. H. Lee, T. Z. Deeb, J. A. Walker, P. A. Davies, S. J. Moss, NMDA receptor activity downregulates KCC2 resulting in depolarizing GABA(A) receptor mediated currents. Nature neuroscience. 14, 736–743 (2011).

8. Y. Ben-Ari, I. Khalilov, K. T. Kahle, E. Cherubini, The GABA Excitatory/Inhibitory Shift in Brain Maturation and Neurological Disorders. The Neuroscientist. 18, 467–486 (2012).

9. G. Sedmak, N. Jovanov-Milošević, M. Puskarjov, M. Ulamec, B. Krušlin, K. Kaila, M. Judaš, Developmental Expression Patterns of KCC2 and Functionally Associated Molecules in the Human Brain. Cereb Cortex. 26, 4574–4589 (2016).

10. T. M. Hyde, B. K. Lipska, T. Ali, S. V. Mathew, A. J. Law, O. E. Metitiri, R. E. Straub, T. Ye, C. Colantuoni, M. M. Herman, L. B. Bigelow, D. R. Weinberger, J. E. Kleinman, Expression of GABA Signaling Molecules KCC2, NKCC1, and GAD1 in Cortical Development and Schizophrenia. J.Neurosci. 31, 11088–11095 (2011).

11. C. Rivera, H. Li, J. Thomas-Crusells, H. Lahtinen, T. Viitanen, A. Nanobashvili, Z. Kokaia, M. S. Airaksinen, J. Voipio, K. Kaila, M. Saarma, BDNF-induced TrkB activation down-regulates the K+–Cl− cotransporter KCC2 and impairs neuronal Cl− extrusion. J Cell Biol. 159, 747–752 (2002).

12. C. Rivera, J. Voipio, J. Thomas-Crusells, H. Li, Z. Emri, S. Sipilä, J. A. Payne, L. Minichiello, M. Saarma, K. Kaila, Mechanism of Activity-Dependent Downregulation of the Neuron-Specific K-Cl Cotransporter KCC2. J. Neurosci. 24, 4683–4691 (2004).

13. W. Löscher, M. A. Rogawski, How theories evolved concerning the mechanism of action of barbiturates. Epilepsia. 53, 12–25 (2012).

14. S. M. Massa, T. Yang, Y. Xie, J. Shi, M. Bilgen, J. N. Joyce, D. Nehama, J. Rajadas, F. M. Longo, Small molecule BDNF mimetics activate TrkB signaling and prevent neuronal degeneration in rodents. J Clin Invest. 120, 1774–1785 (2010).

15. S. K. Kang, G. J. Markowitz, S. T. Kim, M. V. Johnston, S. D. Kadam, Age- and sex-dependent susceptibility to phenobarbital-resistant neonatal seizures: role of chloride co-transporters. Frontiers in Cellular Neuroscience. 9, 173 (2015).

16. J. Shen, K. Ghai, P. Sompol, X. Liu, X. Cao, P. M. Iuvone, K. Ye, N-acetyl serotonin derivatives as potent neuroprotectants for retinas. Proc Natl Acad Sci U S A. 109, 3540–3545 (2012).

17. S.-W. Jang, X. Liu, C. B. Chan, S. A. France, I. Sayeed, W. Tang, X. Lin, G. Xiao, R. Andero, Q. Chang, K. J. Ressler, K. Ye, Deoxygedunin, a Natural Product with Potent Neurotrophic Activity in Mice. PLoS ONE. 5, e11528 (2010).

18. N. A. Setterholm, F. E. McDonald, J. H. Boatright, P. M. Iuvone, Gram-scale, chemoselective synthesis of N-[2-(5-hydroxy-1H-indol-3-yl)ethyl]-2-oxopiperidine-3-carboxamide (HIOC). Tetrahedron Lett. 56, 3413– 3415 (2015).

19. S. D. Croll, N. Y. Ip, R. M. Lindsay, S. J. Wiegand, Expression of BDNF and trkB as a function of age and cognitive performance. Brain Res. 812, 200–208 (1998).

20. F. Rage, M. Silhol, F. Binamé, S. Arancibia, L. Tapia-Arancibia, Effect of aging on the expression of BDNF and TrkB isoforms in rat pituitary. Neurobiol. Aging. 28, 1088–1098 (2007).

21. M. Silhol, V. Bonnichon, F. Rage, L. Tapia-Arancibia, Age-related changes in brain-derived neurotrophic factor and tyrosine kinase receptor isoforms in the hippocampus and hypothalamus in male rats. Neuroscience. 132, 613–624 (2005).

22. M. J. Webster, M. M. Herman, J. E. Kleinman, C. Shannon Weickert, BDNF and trkB mRNA expression in the hippocampus and temporal cortex during the human lifespan. Gene Expr. Patterns. 6, 941–951 (2006).

23. S. C. Kharod, B. M. Carter, S. D. Kadam, Pharmaco-resistant Neonatal Seizures: Critical Mechanistic Insights from a Chemoconvulsant Model. Dev Neurobiol. 78, 1117–1130 (2018).

24. X. Chen, H. Ye, R. Kuruvilla, N. Ramanan, K. W. Scangos, C. Zhang, N. M. Johnson, P. M. England, K. M. Shokat, D. D. Ginty, A chemical-genetic approach to studying neurotrophin signaling. Neuron. 46, 13–21 (2005).

25. P. A. Kipnis, B. J. Sullivan, S. D. Kadam, Sex-Dependent Signaling Pathways Underlying Seizure Susceptibility and the Role of Chloride Cotransporters. Cells. 8, 448 (2019).

26. A. S. Galanopoulou, Dissociated Gender-Specific Effects of Recurrent Seizures on GABA Signaling in CA1 Pyramidal Neurons: Role of GABAA Receptors. J. Neurosci. 28, 1557–1567 (2008).

27. E. J. Calabrese, L. A. Baldwin, U-shaped dose-responses in biology, toxicology, and public health. Annu Rev Public Health. 22, 15–33 (2001).

28. V. I. Dzhala, D. M. Talos, D. A. Sdrulla, A. C. Brumback, G. C. Mathews, T. A. Benke, E. Delpire, F. E. Jensen, K. J. Staley, NKCC1 transporter facilitates seizures in the developing brain. Nature Medicine. 11, 1205 (2005).

29. S. M. Sato, C. S. Woolley, Acute inhibition of neurosteroid estrogen synthesis suppresses status epilepticus in an animal model. Elife. 5 (2016), doi:10.7554/eLife.12917.

30. A. Zayachkivsky, M. J. Lehmkuhle, J. J. Ekstrand, F. E. Dudek, Ischemic injury suppresses hypoxia-induced electrographic seizures and the background EEG in a rat model of perinatal hypoxic-ischemic encephalopathy. J. Neurophysiol. 114, 2753–2763 (2015).

31. J. M. Rennie, L. S. de Vries, M. Blennow, A. Foran, D. K. Shah, V. Livingstone, A. C. van Huffelen, S. R. Mathieson, E. Pavlidis, L. C. Weeke, M. C. Toet, M. Finder, R. M. Pinnamaneni, D. M. Murray, A. C. Ryan, W. P. Marnane, G. B. Boylan, Characterisation of neonatal seizures and their treatment using continuous EEG monitoring: a multicentre experience. Arch. Dis. Child. Fetal Neonatal Ed. (2018), doi:10.1136/archdischild-2018-315624.

32. R. A. Sheldon, C. Sedik, D. M. Ferriero, Strain-related brain injury in neonatal mice subjected to hypoxia– ischemia. Brain Research. 810, 114–122 (1998).

33. R. A. Sheldon, C. Windsor, D. M. Ferriero, Strain-Related Differences in Mouse Neonatal Hypoxia-Ischemia. Dev. Neurosci., 1–7 (2019).

34. S. K. Kang, N. A. Hawkins, J. A. Kearney, C57BL/6J and C57BL/6N substrains differentially influence phenotype severity in the Scn1a+/− mouse model of Dravet syndrome. Epilepsia Open. 4, 164–169 (2019).

35. P. B. de la Tremblaye, S. M. Benoit, S. Schock, H. Plamondon, CRHR1 exacerbates the glial inflammatory response and alters BDNF/TrkB/pCREB signaling in a rat model of global cerebral ischemia: implications for neuroprotection and cognitive recovery. Progress in Neuro-Psychopharmacology and Biological Psychiatry. 79, 234–248 (2017).

36. X. P. He, E. Pan, C. Sciarretta, L. Minichiello, J. O. McNamara, Disruption of TrkB-Mediated Phospholipase C Signaling Inhibits Limbic Epileptogenesis. Journal of Neuroscience. 30, 6188–6196 (2010).

37. R. D. Almeida, B. J. Manadas, C. V. Melo, J. R. Gomes, C. S. Mendes, M. M. Grãos, R. F. Carvalho, A. P. Carvalho, C. B. Duarte, Neuroprotection by BDNF against glutamate-induced apoptotic cell death is mediated by ERK and PI3-kinase pathways. Cell Death and Differentiation. 12, 1329–1343 (2005).

38. F. M. Longo, S. M. Massa, Small-molecule modulation of neurotrophin receptors: a strategy for the treatment of neurological disease. Nature Reviews Drug Discovery. 12, 507–525 (2013).

39. B. L. Hempstead, Brain-Derived Neurotrophic Factor: Three Ligands, Many Actions. Trans. Am. Clin. Climatol. Assoc. 126, 9–19 (2015).

40. L. Subedi, H. Huang, A. Pant, P. M. Westgate, H. S. Bada, J. A. Bauer, P. J. Giannone, T. Sithisarn, Plasma Brain-Derived Neurotrophic Factor Levels in Newborn Infants with Neonatal Abstinence Syndrome. Front. Pediatr. 5 (2017), doi:10.3389/fped.2017.00238.

41. B. Knusel, S. J. Rabin, F. Hefti, D. R. Kaplan, Regulated neurotrophin receptor responsiveness during neuronal migrationand early differentiation. J. Neurosci. 14, 1542–1554 (1994).

42. H. Oh, D. A. Lewis, E. Sibille, The Role of BDNF in Age-Dependent Changes of Excitatory and Inhibitory Synaptic Markers in the Human Prefrontal Cortex. Neuropsychopharmacology. 41, 3080–3091 (2016).

43. K. Deinhardt, M. V. Chao, Trk receptors. Handb Exp Pharmacol. 220, 103–119 (2014).

44. M. V. Chao, Neurotrophins and their receptors: a convergence point for many signalling pathways. Nat. Rev. Neurosci. 4, 299–309 (2003).

45. K. Gottmann, T. Mittmann, V. Lessmann, BDNF signaling in the formation, maturation and plasticity of glutamatergic and GABAergic synapses. Exp Brain Res. 199, 203–234 (2009).

46. E. J. Huang, L. F. Reichardt, Trk receptors: roles in neuronal signal transduction. Annu Rev Biochem. 72, 609– 642 (2003).

47. D. K. Binder, S. D. Croll, C. M. Gall, H. E. Scharfman, BDNF and epilepsy: too much of a good thing? Trends in Neurosciences. 24, 47–53 (2001).

48. J. O. McNamara, H. E. Scharfman, in Jasper’s Basic Mechanisms of the Epilepsies, J. L. Noebels, M. Avoli, M. A. Rogawski, R. W. Olsen, A. V. Delgado-Escueta, Eds. (National Center for Biotechnology Information (US), Bethesda (MD), ed. 4th, 2012; http://www.ncbi.nlm.nih.gov/books/NBK98186/).

49. C. R. Ruiz, J. Shi, M. K. Meffert, Transcript specificity in BDNF-regulated protein synthesis. Neuropharmacology. 76 Pt C, 657–663 (2014).

50. K.-W. Chen, L. Chen, Epigenetic Regulation of BDNF Gene during Development and Diseases. Int J Mol Sci. 18 (2017), doi:10.3390/ijms18030571.

51. K. R. Maynard, J. L. Hill, N. E. Calcaterra, M. E. Palko, A. Kardian, D. Paredes, M. Sukumar, B. D. Adler, D. V. Jimenez, R. J. Schloesser, L. Tessarollo, B. Lu, K. Martinowich, Functional Role of BDNF Production from Unique Promoters in Aggression and Serotonin Signaling. Neuropsychopharmacology. 41, 1943–1955 (2016).

52. B. Riffault, N. Kourdougli, C. Dumon, N. Ferrand, E. Buhler, F. Schaller, C. Chambon, C. Rivera, J.-L. Gaiarsa, C. Porcher, Pro-Brain-Derived Neurotrophic Factor (proBDNF)-Mediated p75NTR Activation Promotes Depolarizing Actions of GABA and Increases Susceptibility to Epileptic Seizures. Cereb Cortex. 28, 510–527 (2018).

53. R. Nardou, S. Yamamoto, G. Chazal, A. Bhar, N. Ferrand, O. Dulac, Y. Ben-Ari, I. Khalilov, Neuronal chloride accumulation and excitatory GABA underlie aggravation of neonatal epileptiform activities by phenobarbital. Brain. 134, 987–1002 (2011).

54. B. Gu, Y. Z. Huang, X.-P. He, R. B. Joshi, W. Jang, J. O. McNamara, A Peptide Uncoupling BDNF Receptor TrkB from Phospholipase Cγ1 Prevents Epilepsy Induced by Status Epilepticus. Neuron. 88, 484–491 (2015).

55. X. P. He, E. Pan, C. Sciarretta, L. Minichiello, J. O. McNamara, Disruption of TrkB-mediated PLC+¦ signaling inhibits limbic epileptogenesis. J Neurosci. 30, 6188–6196 (2010).

56. M. R. Kelley, R. A. Cardarelli, J. L. Smalley, T. A. Ollerhead, P. M. Andrew, N. J. Brandon, T. Z. Deeb, S. J. Moss, Locally Reducing KCC2 Activity in the Hippocampus is Sufficient to Induce Temporal Lobe Epilepsy. EBioMedicine (2018), doi:10.1016/j.ebiom.2018.05.029.

57. P. Q. Duy, W. B. David, K. T. Kahle, Identification of KCC2 Mutations in Human Epilepsy Suggests Strategies for Therapeutic Transporter Modulation. Front. Cell. Neurosci. 13 (2019), doi:10.3389/fncel.2019.00515.

58. K. Fobian, S. Owczarek, C. Budtz, E. Bock, V. Berezin, M. V. Pedersen, Peptides derived from the solvent-exposed loops 3 and 4 of BDNF bind TrkB and p75NTR receptors and stimulate neurite outgrowth and survival. Journal of Neuroscience Research. 88, 1170–1181 (2010).

59. T. A. Gudasheva, P. Povarnina, I. O. Logvinov, T. A. Antipova, S. B. Seredenin, Mimetics of brain-derived neurotrophic factor loops 1 and 4 are active in a model of ischemic stroke in rats. Drug Des Devel Ther. 10, 3545–3553 (2016).

60. A. Gärtner, D. G. Polnau, V. Staiger, C. Sciarretta, L. Minichiello, H. Thoenen, T. Bonhoeffer, M. Korte, Hippocampal long-term potentiation is supported by presynaptic and postsynaptic tyrosine receptor kinase B-mediated phospholipase Cgamma signaling. J. Neurosci. 26, 3496–3504 (2006).

61. L. Minichiello, TrkB signalling pathways in LTP and learning. Nat. Rev. Neurosci. 10, 850–860 (2009).

62. B. H. Han, D. M. Holtzman, BDNF Protects the Neonatal Brain from Hypoxic-Ischemic InjuryIn Vivo via the ERK Pathway. The Journal of Neuroscience. 20, 5775–5781 (2000).

63. R. Roskoski, ERK1/2 MAP kinases: structure, function, and regulation. Pharmacol. Res. 66, 105–143 (2012).

64. M. Ali Shariati, V. Kumar, T. Yang, C. Chakraborty, B. A. Barres, F. M. Longo, Y. J. Liao, A Small Molecule TrkB Neurotrophin Receptor Partial Agonist as Possible Treatment for Experimental Nonarteritic Anterior Ischemic Optic Neuropathy. Curr. Eye Res. 43, 1489–1499 (2018).

65. F. Gu, I. Parada, T. Yang, F. M. Longo, D. A. Prince, Partial TrkB receptor activation suppresses cortical epileptogenesis through actions on parvalbumin interneurons. Neurobiology of Disease. 113, 45–58 (2018).

66. K. B. Nelson, J. K. Grether, L. A. Croen, J. M. Dambrosia, B. F. Dickens, L. L. Jelliffe, R. L. Hansen, T. M. Phillips, Neuropeptides and neurotrophins in neonatal blood of children with autism or mental retardation. Annals of Neurology. 49, 597–606 (2001).

67. S.-J. Tsai, Is autism caused by early hyperactivity of brain-derived neurotrophic factor? Medical Hypotheses. 65, 79–82 (2005).

68. R. L. Bromley, G. Mawer, J. Clayton-Smith, G. A. Baker, Liverpool and Manchester Neurodevelopment Group, Autism spectrum disorders following in utero exposure to antiepileptic drugs. Neurology. 71, 1923– 1924 (2008).

69. R. L. Bromley, G. E. Mawer, M. Briggs, C. Cheyne, J. Clayton-Smith, M. García-Fiñana, R. Kneen, S. B. Lucas, R. Shallcross, G. A. Baker, Liverpool and Manchester Neurodevelopment Group, The prevalence of neurodevelopmental disorders in children prenatally exposed to antiepileptic drugs. J. Neurol. Neurosurg. Psychiatry. 84, 637–643 (2013).

70. G. Williams, J. King, M. Cunningham, M. Stephan, B. Kerr, J. H. Hersh, Fetal valproate syndrome and autism: additional evidence of an association. Dev Med Child Neurol. 43, 202–206 (2001).

71. L. E. F. Almeida, C. D. Roby, B. K. Krueger, Increased BDNF expression in fetal brain in the valproic acid model of autism. Molecular and Cellular Neuroscience. 59, 57–62 (2014).

72. R. Tyzio, R. Nardou, D. C. Ferrari, T. Tsintsadze, A. Shahrokhi, S. Eftekhari, I. Khalilov, V. Tsintsadze, C. Brouchoud, G. Chazal, E. Lemonnier, N. Lozovaya, N. Burnashev, Y. Ben-Ari, Oxytocin-mediated GABA inhibition during delivery attenuates autism pathogenesis in rodent offspring. Science. 343, 675–679 (2014).

73. R. Cloarec, B. Riffault, A. Dufour, H. Rabiei, L.-A. Gouty-Colomer, C. Dumon, D. Guimond, P. Bonifazi, S. Eftekhari, N. Lozovaya, D. C. Ferrari, Y. Ben-Ari, Pyramidal neuron growth and increased hippocampal volume during labor and birth in autism. Science Advances. 5, eaav0394 (2019).

74. T. Nomura, T. F. Musial, J. J. Marshall, Y. Zhu, C. L. Remmers, J. Xu, D. A. Nicholson, A. Contractor, Delayed Maturation of Fast-Spiking Interneurons Is Rectified by Activation of the TrkB Receptor in the Mouse Model of Fragile X Syndrome. J. Neurosci. 37, 11298–11310 (2017).

75. E. E. Zahavi, N. Steinberg, T. Altman, M. Chein, Y. Joshi, T. Gradus-Pery, E. Perlson, The receptor tyrosine kinase TrkB signals without dimerization at the plasma membrane. Sci. Signal. 11, eaao4006 (2018).

76. L. Marchetti, A. Callegari, S. Luin, G. Signore, A. Viegi, F. Beltram, A. Cattaneo, Ligand signature in the membrane dynamics of single TrkA receptor molecules. J. Cell. Sci. 126, 4445–4456 (2013).

77. I. N. Maruyama, Mechanisms of Activation of Receptor Tyrosine Kinases: Monomers or Dimers. Cells. 3, 304–330 (2014).

78. J. Zhai, W. Zhou, J. Li, C. R. Hayworth, L. Zhang, H. Misawa, R. Klein, S. S. Scherer, R. J. Balice-Gordon, R. G. Kalb, The in vivo contribution of motor neuron TrkB receptors to mutant SOD1 motor neuron disease. Hum Mol Genet. 20, 4116–4131 (2011).

79. P. Hu, R. G. Kalb, BDNF heightens the sensitivity of motor neurons to excitotoxic insults through activation of TrkB. Journal of Neurochemistry. 84, 1421–1430 (2003).

80. R. A. Hill, M. van den Buuse, Sex-dependent and region-specific changes in TrkB signaling in BDNF heterozygous mice. Brain Res. 1384, 51–60 (2011).

81. H. E. Scharfman, N. J. MacLusky, Differential regulation of BDNF, synaptic plasticity and sprouting in the hippocampal mossy fiber pathway of male and female rats. Neuropharmacology. 76 (2014), doi:10.1016/j.neuropharm.2013.04.029.

82. H. E. Scharfman, N. J. Maclusky, Similarities between actions of estrogen and BDNF in the hippocampus: coincidence or clue? Trends Neurosci. 28, 79–85 (2005).

83. A. S. Galanopoulou, A. Kyrozis, O. I. Claudio, P. K. Stanton, S. L. Moshé, Sex-specific KCC2 expression and GABA(A) receptor function in rat substantia nigra. Exp. Neurol. 183, 628–637 (2003).

84. A. S. Galanopoulou, Sex- and cell-type-specific patterns of GABAA receptor and estradiol-mediated signaling in the immature rat substantia nigra. European Journal of Neuroscience. 23, 2423–2430 (2006).

85. T. S. Perrot-Sinal, C. J. Sinal, J. C. Reader, D. B. Speert, M. M. McCarthy, Sex differences in the chloride cotransporters, NKCC1 and KCC2, in the developing hypothalamus. J. Neuroendocrinol. 19, 302–308 (2007).

