## Supplementary figures and images for "TrkB agonists prevent post-ischemic BDNF-TrkB mediated emergence of refractory neonatal seizures in CD-1 pups"

### Graphical abstract

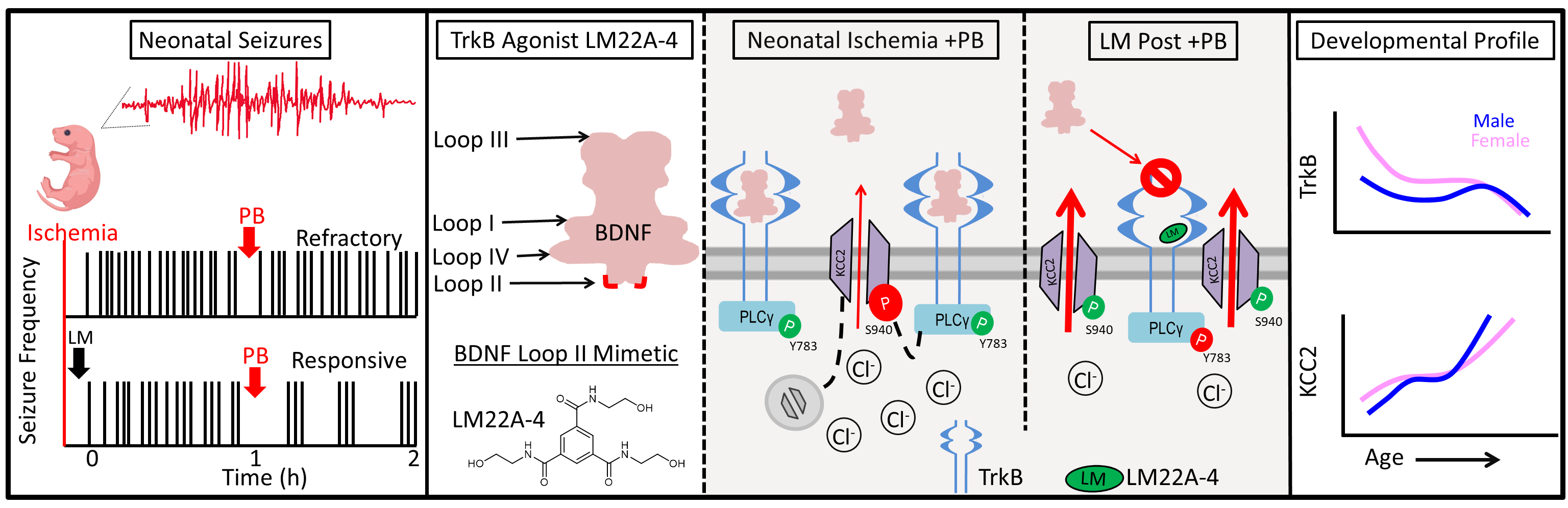
